# Multiplex Base Editing to Protect from CD33-Directed Therapy: Implications for Immune and Gene Therapy

**DOI:** 10.1101/2023.02.23.529353

**Authors:** Florence Borot, Olivier Humbert, Gregory A Newby, Emily Fields, Sajeev Kohli, Stefan Radtke, George S. Laszlo, Thiyagaraj Mayuranathan, Abdullah Mahmood Ali, Mitchell J. Weiss, Jonathan S. Yen, Roland B. Walter, David R. Liu, Siddhartha Mukherjee, Hans-Peter Kiem

**Author notes:** These authors contributed equally (Florence Borot, Olivier Humbert, Siddhartha Mukherjee, Hans-Peter Kiem).

## Abstract

On-target toxicity to normal cells is a major safety concern with targeted immune and gene therapies. Here, we developed a base editing (BE) approach exploiting a naturally occurring CD33 single nucleotide polymorphism leading to removal of full-length CD33 surface expression on edited cells. CD33 editing in human and nonhuman primate (NHP) hematopoietic stem and progenitor cells (HSPCs) protects from CD33-targeted therapeutics without affecting normal hematopoiesis *in vivo*, thus demonstrating potential for novel immunotherapies with reduced off-leukemia toxicity. For broader applications to gene therapies, we demonstrated highly efficient (>70%) multiplexed adenine base editing of the CD33 and gamma globin genes, resulting in long-term persistence of dual gene-edited cells with HbF reactivation in NHPs. *In vitro*, dual gene-edited cells could be enriched via treatment with the CD33 antibody-drug conjugate, gemtuzumab ozogamicin (GO). Together, our results highlight the potential of adenine base editors for improved immune and gene therapies.

**Graphical abstract:** 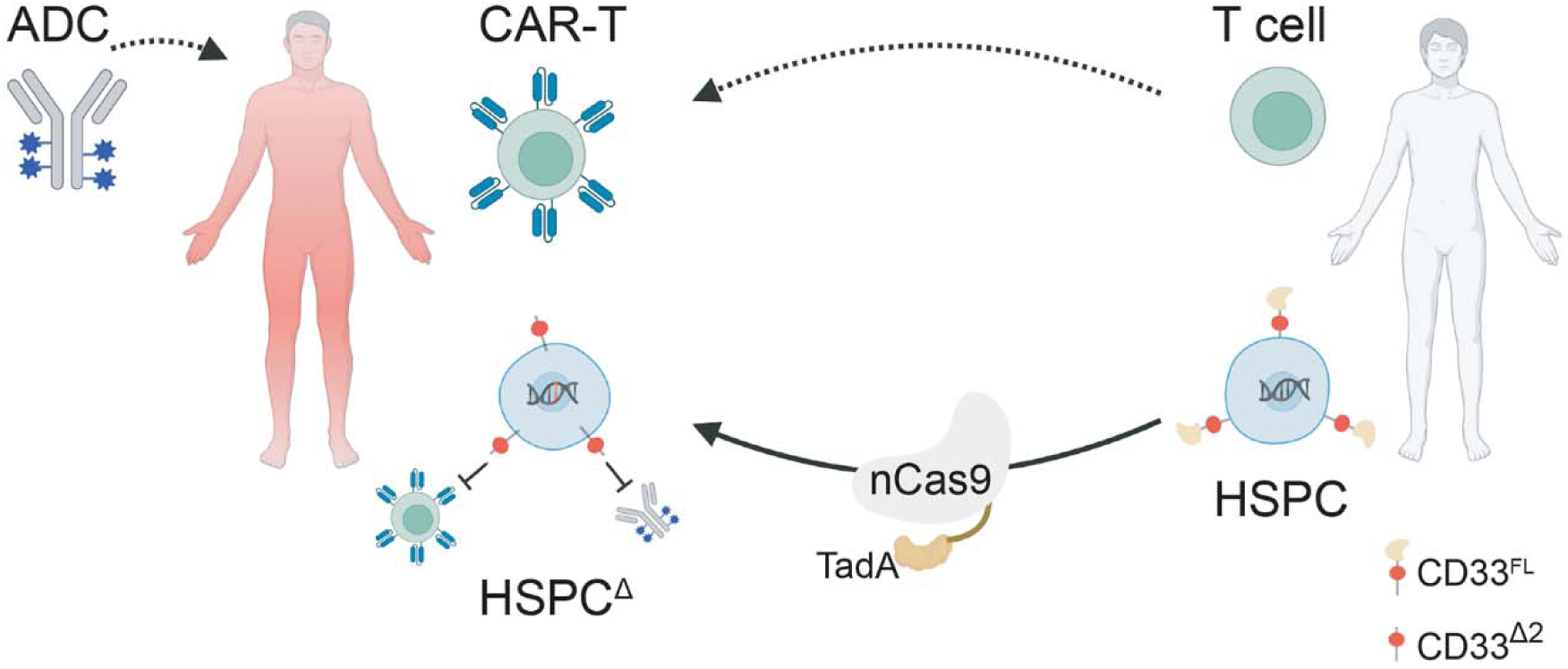

## Introduction

There is increasing interest in immunotherapeutic strategies to treat cancer, with a growing list of drugs approved for clinical use. Because of the lack of cancer-specific antigens as suitable targets for immune and gene therapies in many instances, on-target toxicity to normal cells is a major safety concern with targeted immune and gene therapies. For example, with treatments for hematologic malignancies, expression of target antigens on normal hematopoietic cells can lead to prolonged, severe myelosuppression and associated adverse sequelae such as hypogammaglobulinemia and/or life-threatening/limiting bleeding or infection. For immunotherapy directed at CD33, a myeloid differentiation antigen expressed broadly on neoplastic cells in patients with acute myeloid leukemia (AML), we and others proposed to create a specific leukemia antigen by using CRISPR/Cas9 to engineer CD33-negative normal donor hematopoietic stem and progenitor cells (HSPCs) ^1–3^. Ablation of CD33 on HSPCs did not alter functionality or hematopoietic repopulation capacity and conferred resistance to CD33-targeted immunotherapies like the clinically approved CD33 antibody-drug conjugate gemtuzumab ozogamicin (GO) or CD33-directed chimeric antigen receptor (CAR) T cells *in vitro* and *in vivo*. CRISPR/Cas9 has proven very efficient at editing genes, but its double strand break (DSB)-based mechanism has been associated with adverse consequences such as p53 activation, large insertions/deletions, and chromosomal translocations in some cell types including HSPCs^4–7^. While the off-target assessment of our previous study did not uncover deleterious mutations in CRISPR/Cas9-edited HSPCs, the long-term effects of a lineage marker knockout in hematopoietic cells remain uncertain.

Gene editing of normal HSPCs to confer immunotherapy resistance could be harnessed as an enrichment and selection strategy for cells simultaneously edited at another locus to treat genetic or infectious diseases. Such enrichment might overcome a current limitation in the field of HSPC gene therapy, namely the inability to achieve a high proportion of gene-modified cells in patients. *In vivo* selection of gene-modified cells might also enable the use of conditioning with reduced toxicity such as nonmyeloablative^8^ and nongenotoxic regimens^9, 10^.

Previous strategies to enrich gene-modified HSPCs include overexpression of mutated O6-methylguanine-DNA methyltransferase (MGMT), conferring resistance to nitrosoureas and O6-benzylguanine drugs^11, 12^. Stable HSPC enrichment could be achieved with this approach, but safety has remained a concern due to the use of lentiviral resistance gene expression and need for alkylating agents for selection. Introduction of genes that confer resistance to a drug or small molecule^13, 14^ has also been investigated for selecting cells with a therapeutic gene modification. However, translation of these approaches to HSPC-based therapies has been challenging given they might negatively affect the cells’ fitness and multilineage engraftment capacity. Additional investigation is thus needed for the development of *in vivo* selection strategies for HSPCs.

Base Editors (BEs) can precisely introduce single nucleotide mutations through a DSB-independent mechanism to either rescue gene expression or generate nonpathogenic gene variants^15–18^. Here we investigated its potential therapeutic applications to: 1) engineer a healthy donor allogeneic HSPC transplant product resistant to CD33-targeted therapies, and 2) develop a multiplex base editing strategy for selection of a therapeutic edit in the gamma globin genes (HBG) that increases fetal hemoglobin (HbF) expression by modulating the binding of key regulatory factors^19–21^, including the repressor protein BCL11A^22^.

Here, we harnessed the inherent properties of BEs^23–26^ by specifically employing the adenine BE ABE8e^23^ to target CD33 in HSPCs alone and in combination with an edit in the *HBG1/2* promoters. Using mouse xenograft and NHP transplantation models, we demonstrate that these cells show unaltered long-term engraftment and multi-lineage hematopoietic reconstitution with resistance to CD33-directed drugs *in vitro* and *in vivo*. Together, these results highlight the potential of BE for cancer immunotherapies and more generally for HSPC gene therapies.

## Results

### ABE8e editing of CD33 exon2 splicing acceptor site leads to exon 2 skipping

The dominant full-length isoform of CD33 (CD33^FL^) is composed of 7 exons. Most current anti-CD33 investigational treatments including GO are directed against epitopes localized in the V-set domain of CD33 encoded by exon 2. The single nucleotide polymorphism rs12459419, located within the exon splicing enhancer (ESE) site of exon 2, substitutes C for T (changing Alanine 14 to Valine, A14V), resulting in the increased expression of a shorter isoform (CD33^Δ2^) and reduced translation of CD33^FL^ ^27^. Exome and genome sequencing databases revealed that 30% in all population is homozygous TT for this SNP^28, 29^. Splicing quantitative trait loci (sQTL) analysis of the genotype-tissue expression (GTEx) database indicated genotype dependent excision of exon 2^30, 31^. The investigation of the effect of this splicing polymorphism^32–35^ on pediatric AML patient response to GO treatment revealed that the CC genotype is associated with greater CD33 expression on AML blasts, as well as lower risk of relapse and higher survival.

To generate a cell population resistant to cancer immunotherapies targeting CD33 exon 2, we assessed whether cytosine base editors (CBEs) and ABEs^24, 36, 37^ could edit the CD33 ESE site and exon 2 acceptor splice site to reduce the expression of CD33^FL^. Owing to the properties of BEs^24, 36, 37^, we devised two strategies: either by replicating the known rs12459419 SNP using the CBE BE4max, or by mutating the exon 2 splicing acceptor site using ABE8e (Fig. 1a-c). Two sgRNAs were tested with BE4max protein: sgCBE_1_ with an editing window targeting the ESE, and sgCBE_2_ which could concomitantly edit the splicing acceptor and the ESE of CD33 exon2 (Fig. 1c). Both sgRNAs led to more than 50% editing at the targeted nucleotides, but also produced indels as shown by Sanger sequencing and high throughput sequencing (HTS) (Fig. 1d-e) 6 to 7 days post electroporation. The BE4max edited cells displayed approximately 65% to 69% loss of CD33 expression when assessed by flow cytometry after staining with the P67.6 CD33 antibody, which recognizes an epitope encoded withing exon 2 and is used as the targeting component for GO therapy. Interestingly, a ribonucleoprotein complex made of ABE8e protein and sgABE led to more than 95% editing at the targeted splicing acceptor adenine and almost no detectable indels. A bystander adenine (a_1_) localized within intron 1 was also edited (Fig. 1d-e). The efficient conversion of the exon 2 splicing acceptor adenine results in more than 94% of HSPCs that were not recognized by the anti-CD33 clone P67.6 (Fig. 1f). To assess the outcomes of the CBE and ABE editing strategies, we harvested the total RNA of unedited or edited HSPCs and performed RT-PCR with different sets of primers to detect CD33^FL^ and/or CD33^Δ2^ cDNAs. As a control, we used CD34^+^ unedited cells that were homozygous for CC at the A14V SNP and expressed mostly CD33^FL^ mRNA. Separation of the PCR products by gel electrophoresis (Fig1g) displays clear exon 2 skipped amplicons (highlighted with red and green stars) in all edited cells. Sanger sequencing confirmed the sequence of ABE-generated CD33^Δ2^ cDNA. We also noted the presence of CD33^FL^ cDNA (highlighted with blue and black stars) in CBE edited HSPCs, which correlates with the detection of 30% CD33 expression by FACS. Finally, staining of ML-1 cells with a set of CD33 antibodies^38^, either specific to the V set domain (P67.6), or to the C2 domain of all CD33 isoforms (9G2), or to the C2 domain only when exon 2 is absent (11D5), showed that CD33 expression at the membrane is lost following ABE8e editing (Fig. 1h). Collectively these observations highlight the feasibility and efficacy to induce exon skipping in HSPCs by transient exposure to ABE8e/sgABE RNP mediated editing.

**Fig. 1.**
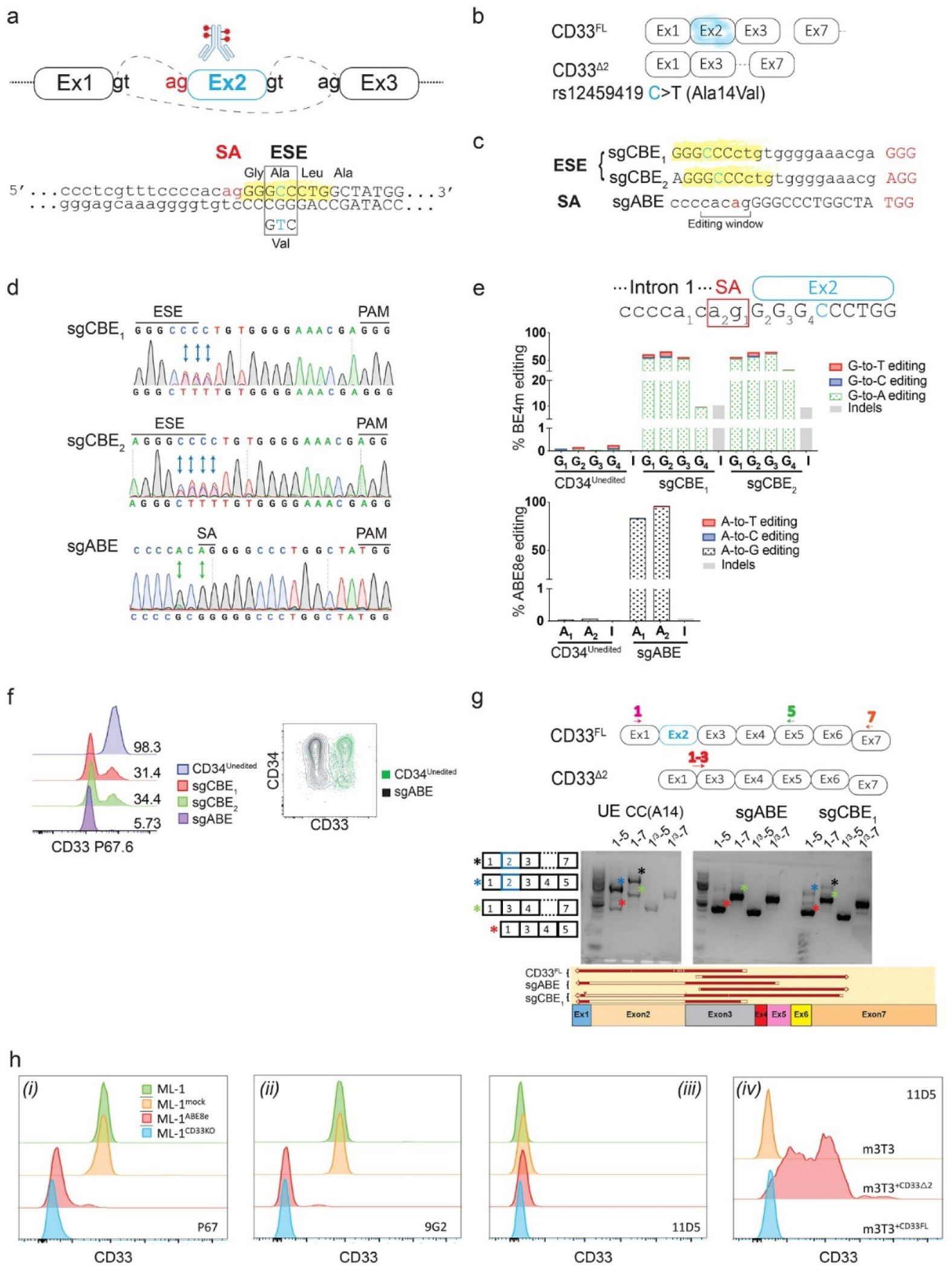
ABE8e introduces A>G conversion at targeted nucleotide with up to 95% efficiency and negligeable indels. a, *Top* Details of exon 1-3 region and possible splicing outcomes, ag: Splicing acceptor site (SA). gt: Splicing donor site. Dotted lines depict possible splicing event. GO Gentuzumab Ozogamicin (Ab icon) recognizes an epitope located in exon 2. *Bottom* Details of the Intron1(lower case)/exon2(upper case) junction DNA sequence with highlight of exon 2 SA (red) and Exon Splicing Enhancer site (ESE in yellow). b, CD33 mRNA full length (CD33FL) contains 7 exons, exon 2 encodes an Ig-like V-type domain. CD33^Δ2^ lacks exon 2 due to a common polymorphism (rs12459419) which changes C>T resulting in an altered exonic splicing enhancer (ESE) site. c, Sequences of the protospacers design to either edit the ESE with BE4max (sgCB_1_ and CB_2_) or the SA with ABE8e (sgABE). PAM is in red. d, Sanger sequencing profiles of edited cells are compared to the wild-type genomic sequence (top). Editing of a cytosine or an adenine that mutates the ESE or SA is indicated by arrows. e, HTS analysis by CRISPResso2 depicts the editing efficiency at the targeted nucleotides and bystander’s cytidine or adenine, as well as indels. f, FACS analysis of the edited cells 7 days post electroporation with antibody clone P67.6 which recognize an epitope located in exon 2. After BE4-mediated editing, around 30% of the CD34^+^cells show CD33 expression while less than 5% after ABE-mediated editing. g, ESE or SA editing induces exon 2 skipping. Editing outcomes assessment. Exon skipping in CD34^+^ edited cells was characterized by performing PCR on cDNA with sets of primers, specific to CD33^Δ2^ (spanning exon junction 1-3), or common to all isoforms (in exons 1, 5 and 7). PCR products were separated by polyacrylamide gel electrophoresis and visualized by SYBR-safe fluorescence. Sanger sequencing of PCR products confirm the absence of exon 2 in edited cells while all other exons are intact. h, ABE8e modification of CD33 locus in ML-1 cells results in a loss of CD33 cell surface expression. ML-1 cells were mock electroporated (ML-1^mock^), or electroporated with ABE8e and CD33 monoclonal antibody (mAb) staining compared to parental ML-1 cells or ML-1 cells in which both alleles of CD33 had been disrupted via CRISPR targeting of exon 1 (ML-1^CD33KO^). P67 mAb *(i)* binds to the V-set domain of CD33, 9G2 *(ii)* binds to the C2-set domain of CD33, whether V-set is present or not, and 11D5 (*iii* and *iv*) binds to the C2-set domain of CD33 when the V-set is absent, e.g., CD33 lacking exon 2. Specificity of 11D5 to CD33^△2^ is shown *(iv.)* using mouse 3T3 cells that lack human CD33 expression are shown (m3T3) with forced expression of either CD33^△2^ (m3T3^+CD^^33△2^) or full length CD33 (m3T3^+CD33FL^

### CD33^Δ2^ cells display intact phagocytic function and resistance to CD33 targeted therapy *in vitro*

To assess the functional capacity of CD33^Δ2^ edited hematopoietic cells, we evaluated their phagocytotic ability by incubating *in vitro* myeloid differentiated unedited or CD33^Δ2^-edited CD34^+^ cells with *E. coli* bioparticles (Fig. 2a). The quantification of internalization of *E. coli* bioparticles by flow cytometry showed similar level of phagocytosis between edited and unedited cells. Furthermore, the inhibition of actin polymerization with cytochalasin D blocked the phagocytic capacity of unedited and CD33^Δ2^-edited myeloid cells similarly.

**Fig. 2.**
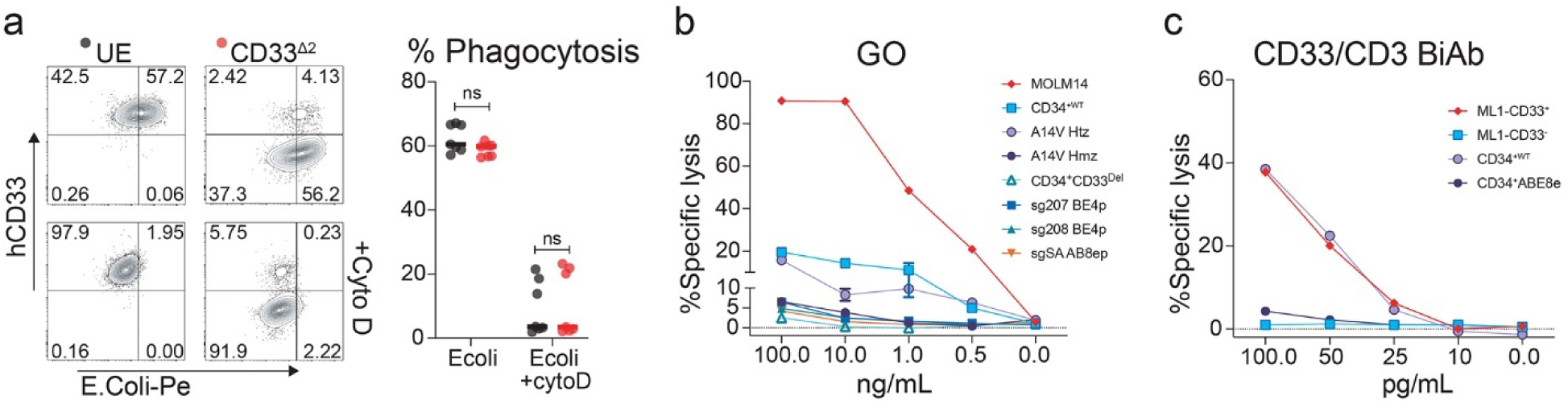
ABE8e edited HSPCs are resistant to CD33-targeted therapy *in vitro* and *in vitro* differentiated CD33^Δ2^-edited myeloid cells display intact phagocytic capacity. a, *In vitro* differentiated unedited (UE) or CD33^Δ2^-edited monocytes show comparable phagocytosis capacity, as measured by *E.Coli* bioparticles internalization. *Left,* Representative FACs plots of E.Coli bioparticles internalization. Treatment with actin polymerization inhibitor, cytochalasin D abrogates phagocytosis. *Right,* Graph of phagocytosis quantification. Unpaired t test. b, CD33^Δ2^-edited CD34^+^ cells resist GO cytotoxicity *in vitro*. Cells were incubated 48 hours with GO and cytotoxicity analyzed by FACS using Sytox Blue or 7AAD as a viability dye. CD33^Δ2^-edited CD34^+^ show same resistance to GO cytotoxicity than a donor with homozygous rs12459419 A14V SNP. (2 independent experiments, 2 donors). c, ML1 CD33 WT or KO cells and non-edited (WT), or ABE8e-edited mPB CD34^+^ cells were assessed for resistance to the CD33/CD3 bispecific T-cell engager (generated from published sequences and described in Correnti et al.^60^). Target cells were incubated with healthy donor T cells for 2 days and absolute cell number and viability were detected by flow cytometry analysis following staining with 4’,6-diamidino-2-phenylindole (DAPI). Results were normalized to reactions that were not treated with FH330 (1 independent experiment run in triplicates, 1 donor).

Additionally, we subjected ABE8e-edited CD34^+^ CD33^Δ2^ cells to GO to evaluate whether editing confers resistance to CD33-targeted therapy. While CD34^+ Unedited^ cells predominantly expressing CD33^FL^ display high sensitivity to GO, CD33^Δ2^-edited CD34^+^ cells show GO resistance, similar to a control donor cells homozygous for CC genotype (A14) and to CD33 knockout CD34^+^ cells (CD34^+^CD33^Del^). MOLM14 leukemic cells, homozygous for CC genotype (A14) and expressing a high level of CD33, were also used as controls, showing high sensitivity to GO (Fig. 2b and Fig. S1). Furthermore, CD34^+^ cells edited with ABE8e were almost completely resistant to the cytotoxic activity of a CD33/CD3 bispecific T cell engager that is dependent on an exon 2 encoded epitope (Fig. 2c), thus corroborating results from GO treatment. Taken together, these results confirm that CD34^+^CD33^Δ2^ base edited cells retain intact phagocytosis capacity and are insensitive to CD33 targeted therapies *in vitro*.

### Edited HSPCs engraft and repopulate a complete hematopoietic system

Next, we examined if adenine base editing of CD34^+^ cells could perturb hematopoiesis by following the engraftment and hematopoietic repopulation of CD34^+ Unedited^ and CD34^+^ CD33^Δ2^ cells in mice over time (Fig3.a). In all three analyzed tissues (peripheral blood, bone marrow, spleen) by flow cytometry, staining of the engrafted cells with the P67.6 antibody clone barely detected CD33 expression in mice reconstituted with CD33^Δ2^ -edited CD34^+^cells compared to CD34^+ Unedited^ controls (Fig. 3b and c) at 16 weeks post-transplantation. Flow cytometry analysis of mice transplanted with control CD34^+Unedited^ or CD33^Δ2^ -edited CD34^+^ cells revealed no differences in the levels of overall human cell engraftment (hCD45^+^) or of specific donor-derived hematopoietic lineages (Fig. 3a and c, Fig. S2). High throughput sequencing analysis of ABE8e-edited human donor cells pre-and post-transplantation showed that editing efficiency is maintained *in vivo* (Fig3d. Furthermore, similar bone marrow architecture was observed in both groups engrafted with unedited or CD33^Δ2^-edited cells as illustrated by H&E staining (Fig. S3).

**Fig. 3.**
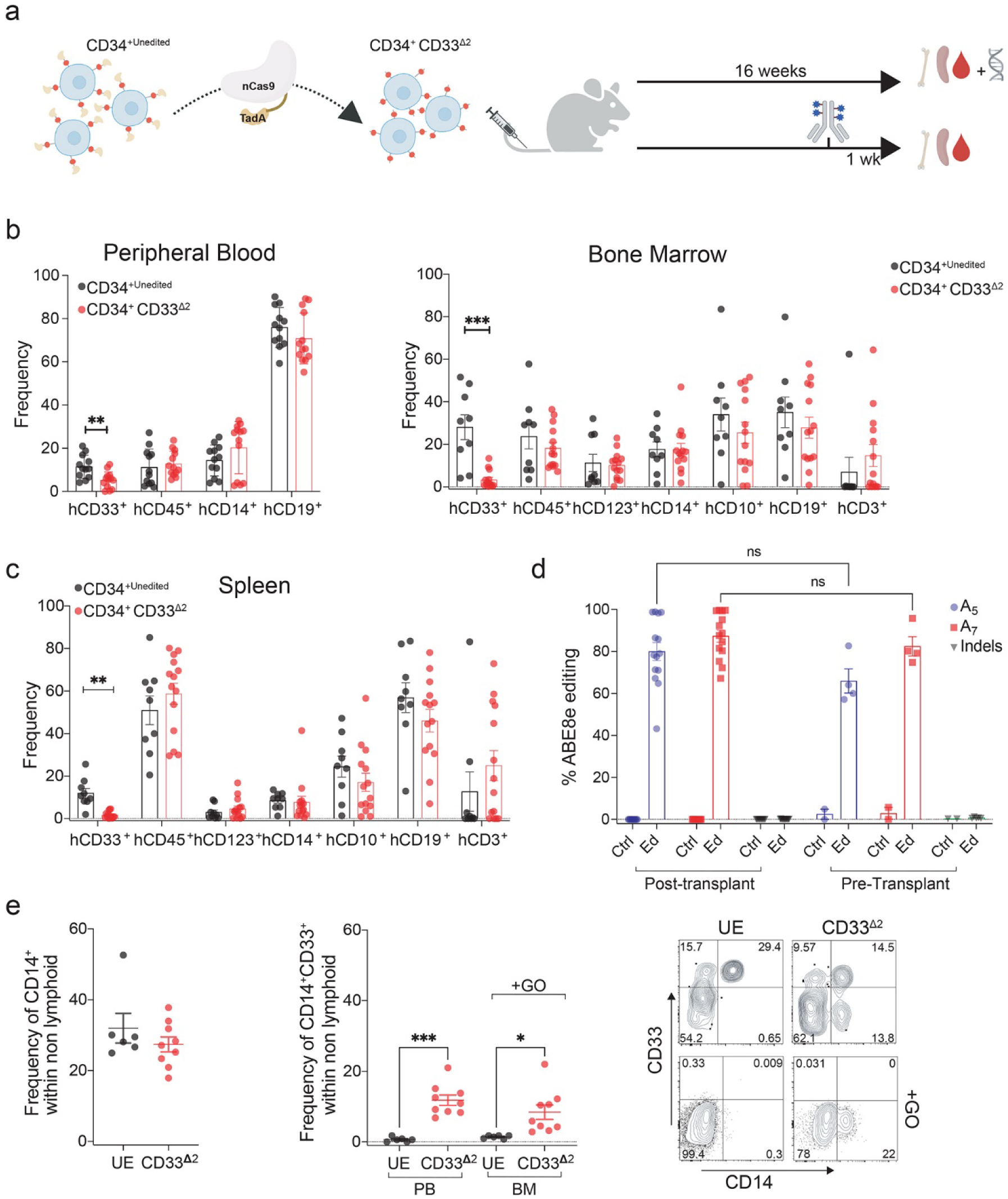
CD33^Δ2^-edited CD34^+^ engraft, recapitulate a complete hematopoietic system and display resistance to GO CD33-targeted therapy *in vivo.* a, Schematic of the experiment b, c, Measure of engraftment by percentage of human CD45^+^ cells and of hematopoietic repopulation by frequency of progenitors myeloid (CD123) and lymphoid (CD10), as well as mature myeloid (CD14) and lymphoid (CD19), and T cells (CD3) within the human CD45 population in peripheral blood at 8 weeks post-transplantation and in the bone marrow and the spleen at 16 weeks post-transplantation. d, On-target editing (HTS analysis) in edited cells kept *in vitro* (pre-Transplant) or harvested in the BM 16 weeks post-Transplantation. e, *Left*, frequency of CD14^+^ myeloid cells in the PB of 12 weeks post-transplanted mice. *Middle*, frequency of CD14^+^CD33^-^ cells in the PB of 12 weeks post-transplanted mice before GO treatment and in the BM one week after GO treatment (2.5 ug per mouse). *Right*, representative flow plots show eradication of CD14^+^ cells in control group (Ctrl, unedited cells) while CD14^+^ cells are detected in BM of mice engrafted with CD33^Δ2^-edited CD34^+^ cells. (UE, unedited cells) (2 independent experiments, 2 donors). One way ANOVA.

To determine whether ABE8e-editing of CD33 exon 2 splice site rendered engrafted CD33^Δ2^-edited cells resistant to targeted immunotherapy *in vivo*, CD34^+ Unedited^ and CD33^Δ2^-edited CD34^+^ transplanted mice were injected with GO (Fig3.a). In both mouse groups, prior to GO treatment, the frequencies of CD14^+^ cells at 12 weeks post-transplantation were comparable (Fig. 3e, left panel). However, following GO treatment, the CD33^-^CD14^+^ population was present only in CD33^Δ2^-edited CD34^+^injected mice (Fig. 3e, center and right panels). Remarkably, while a single dose of GO resulted in eradication of the CD33^+^ population, the CD33^-^CD14^+^ population remained detectable at a significant frequency in mice transplanted with edited cells (Fig. 3e, Fig. S4).

Together, these data indicate that ABE-mediated CD33 exon 2 skipping does not impair bone marrow reconstitution or hematopoietic differentiation of CD34^+^ HSCs and confers resistance to CD33 targeted therapy.

### CIRCLE seq and *in vivo* off target analysis

To characterize off-target editing resulting from base editor treatment targeted to CD33, we conducted CIRCLE-seq^39^ to experimentally nominate off-target sites. CIRCLE-seq nominated a list of 754 potential off-target sites for sgABE and 548 sites for sgCBE_1_. We ranked these sites based on their frequency of occurrence and chose the top 20 nominated sites from each list for further characterization by HTS in ABE edited CD34^+^ cells (Fig. 4 and 5, Fig. S5).

**Fig. 4.**
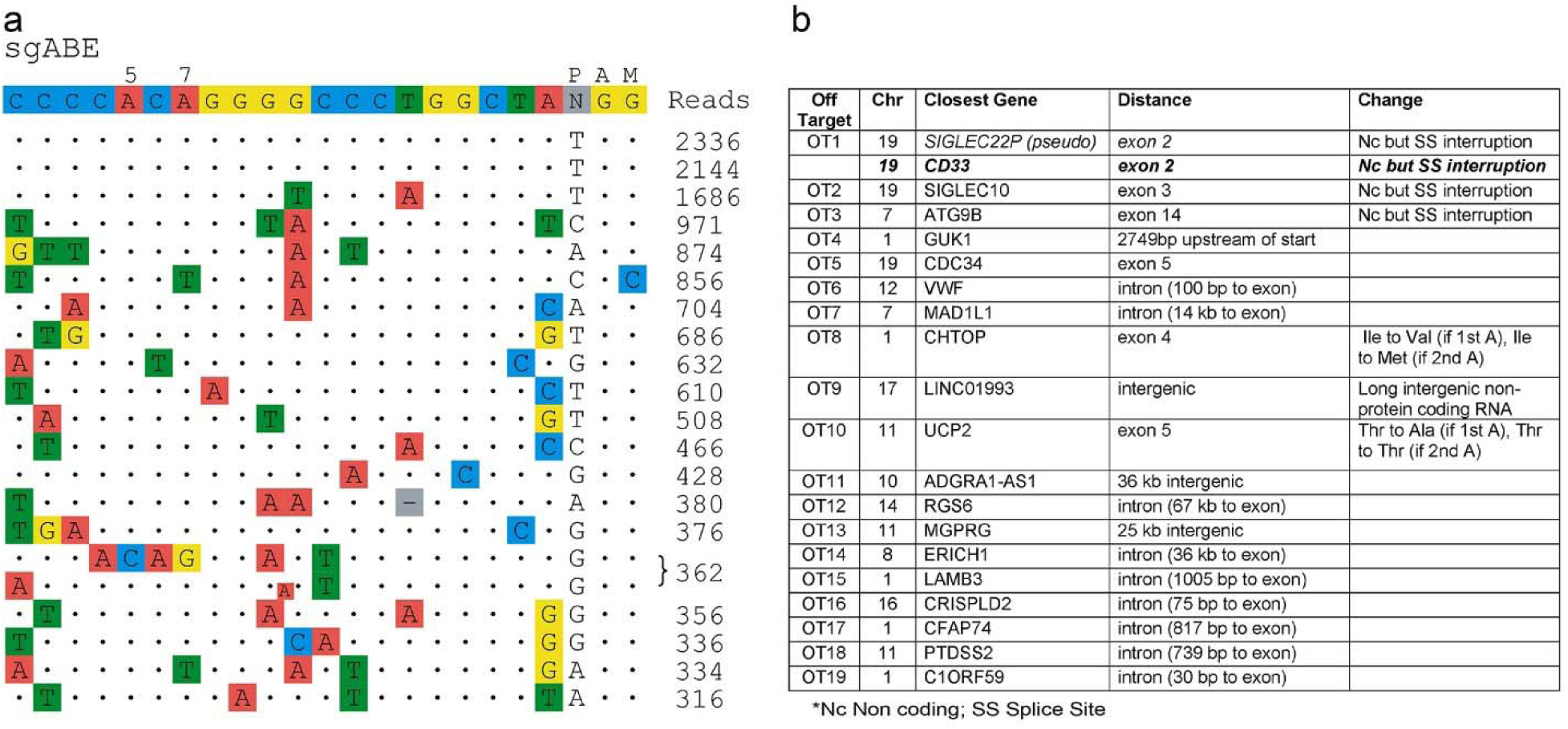
Off target associated with ABE8e protein editing of CD33 exon 2 SA in CD34^+^ cells. a, Visualization of on target and top 19 off target sites detected by CIRCLE-seq. Read counts and alignment of each site to the sgABE protospacer is shown. b, Table summarizing the 19 top off target identified loci.

**Fig. 5.**
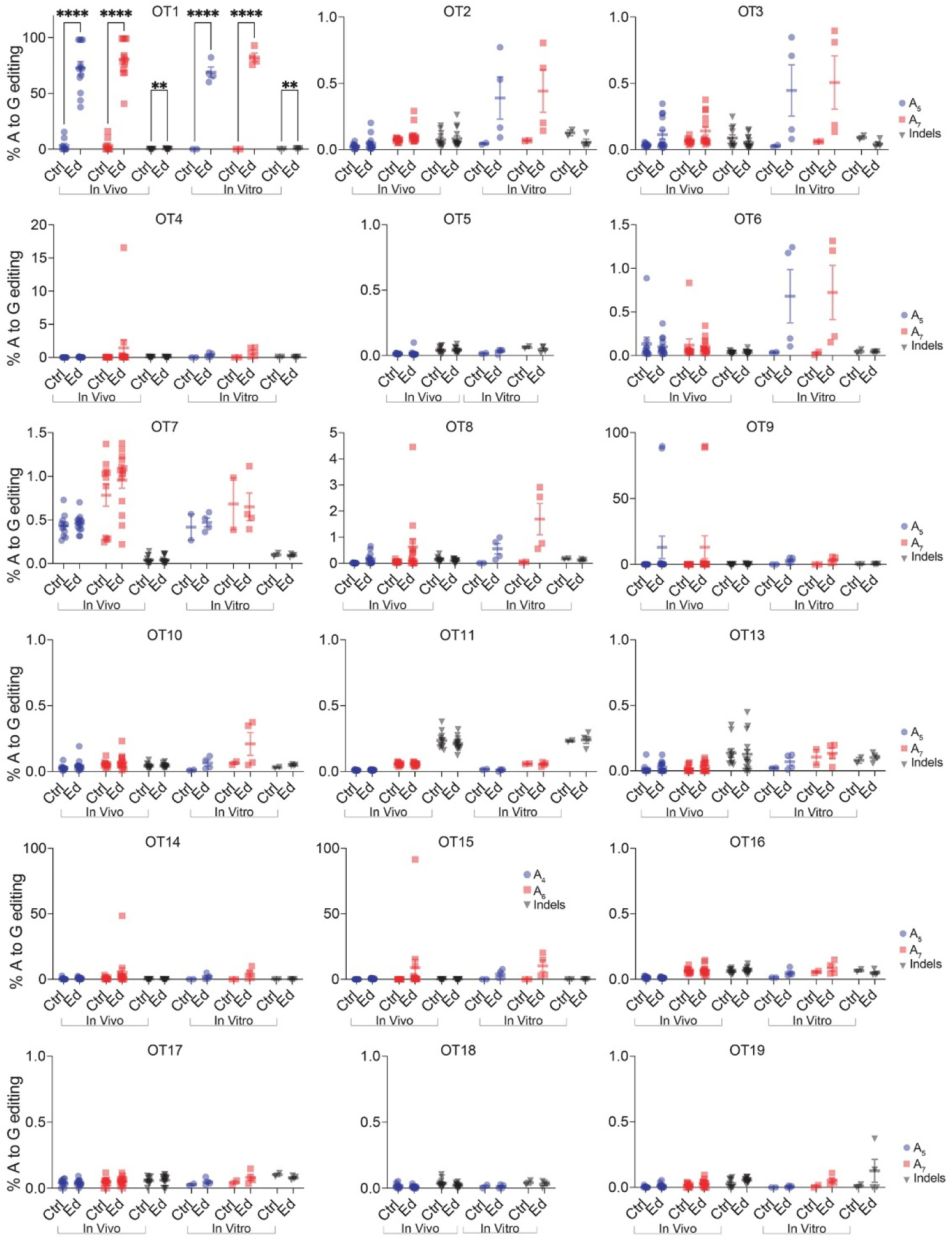
Quantification and assessment of Off target associated with ABE8e protein editing of CD33 exon 2 SA in unedited or CD33^Δ2^ engrafted cells. Editing assessment of A-to-G editing at position A5, A7 and indels at the 19 identified top off target loci in cells kept in vitro or in engrafting human unedited (control, Ctrl) or edited (Ed) cells from bone marrow at 16 weeks post-transplantation (noted, *In Vivo*). Sixteen weeks after transplantation, the top 19 human off-target loci identified by CIRCLE-seq were amplified from genomic DNA from the bone marrow of mice. Amplicons were sequenced by HTS and A-to-G editing at position A5 and A7 was quantified. In vivo analysis from 2 donors, 2 independent experiments; In vitro analysis from 2 donors, 4 independent experiments. Unpaired t test.

Quantification of off-target ABE editing was assessed by high-throughput Illumina MiSeq sequencing (HTS) in edited CD34^+^ HSPCs and unedited controls 7 days following electroporation with ABE-gRNA RNP complex and maintenance in cell culture. Additionally, to assess editing in engrafting hematopoietic stem cells, we conducted HTS at each site in engrafted cells harvested at 16 weeks post transplantation in mice. Of the 19 off-target loci assessed for sgABE, the majority (13 loci) fell in intergenic or intronic regions, one of which, OT12, was not amenable to sequencing, 3 loci (OT5, OT8, OT10) were in exons, and 3 loci (OT1, OT2, OT3) were in the non-coding region of a splice site (Fig. 4 and 5).

To characterize off-target editing resulting from base editor treatment targeted to CD33, we conducted CIRCLE-seq^39^ to experimentally nominate off-target sites. CIRCLE-seq nominated a list of 754 potential off-target sites for sgABE and 548 sites for sgCBE_1_. We ranked these sites based on their frequency of occurrence and chose the top 20 nominated sites from each list for further characterization, which included the targeted site in CD33 and 19 potential off-target loci each (Fig. 4 and 5, Fig. S5).

Quantification of off-target ABE editing was assessed by high-throughput Illumina MiSeq sequencing (HTS) in edited CD34^+^ HSPCs and unedited controls 7 days following editor electroporation and maintenance in cell culture. Additionally, to assess editing in engrafting hematopoietic stem cells, we conducted HTS at each site in engrafted cells harvested at 16 weeks post transplantation in mice. Of the 19 off-target loci assessed for sgABE, the majority (13 loci) fell in intergenic or intronic regions, one of which, OT12, was not amenable to sequencing, 3 loci (OT5, OT8, OT10) were in exons, and 3 loci (OT1, OT2, OT3) were in the non-coding region of a splice site (Fig. 4 and 5).

Of the three sgABE off-target loci that overlap splice sites, OT1 is a CD33 pseudogene that has 100% identity to the targeted site and was efficiently edited in all treated cells as expected. The off-target editing at OT2 and OT3 is lower than 1% in all samples and the frequency of editing in these sites was slightly reduced in engrafted cells. In OT3, the edited splice site falls after the stop codon of the gene and therefore should not impact the coding sequence.

Of the three tested off-target loci that overlap exons, editing was observed at OT8 with up to 4.45% editing and at OT10 with up to 0.37% editing. Editing at OT8 could lead to two possible amino acid changes in CHTOP: Ile92Met (rs757148531), which occurs naturally and is not associated with any abnormal phenotype^28, 40^ or Ile92Val, predicted to be benign^41^. Editing at OT10 can result in a T126A mutation in UCP2, though a silent edit was more efficient. Further characterization or minimization of these off-target editing sites should be conducted before the application of this method in the clinic.

At some sgABE off-target loci such as OT4, OT9, OT14, and OT15, cells that engrafted in one or two mice were quantified to have high frequencies of off-target editing (15-90%, Fig. 4c). However, the cell lineages and the total human engraftment in these mice were not different than unedited controls, so the increase in observed editing efficiency was not caused by oncogenic clonal expansion and may instead be due to the bottleneck of a small number of edited cells that engrafted durably in mice. We observed that indels resulting from ABE treatment were minimal at all sites including those that were efficiently base edited (Fig. 4 and 5).

Though editing using BE4max was less efficient, we examined its off-target landscape in edited cells maintained *in vitro* (Fig. S5). Of the 19 sgCBE_1_ off-target loci analyzed, the top 3 were observed to be edited following CBE treatment. These 3 edited sites include the homologous splice site CD33 pseudogene, an exon of *SIGLEC25P*, and one in intergenic space. The high frequency of indels observed in the pseudogene and CD33 by CBE carries risks of genomic translocations, which we expect will render CBE editing a less safe strategy in comparison to our ABE approach.

### CD33/HBG multiplex ABE8e treatment of human CD34^+^ HSPCs results in a high frequency of dual-edited cells

Having demonstrated that CD33-edited cells are protected from CD33 drug cytotoxicity, we sought to determine whether a similar strategy could be exploited to enrich for an edit that is therapeutically relevant for genetic diseases such as hemoglobinopathies, when the CD33 edit and the therapeutic edit are installed simultaneously in cells. As a therapeutic model system, we focused on editing the regulatory region of HBG genes, which has previously been targeted by site-directed nucleases^19, 21, 42, 43^ or by BE^44, 45^ to reactivate the expression of γ-globin and fetal hemoglobin (α2γ2) as treatment for beta-hemoglobinopathies. We specifically chose to target the HBG promoters at position -175 relative to the initiation codon, where a natural T-to-C mutation results in hereditary persistence of fetal hemoglobin by creating a de novo binding site for the transcriptional activator TAL1,, ultimately increasing HBG expression^46–48^. BE targets for this HBG-175 site have previously been investigated and validated for HbF reactivation^45, 49^. For optimal targeting of this HBG site, we use the ABE8e-NG variant in which the Cas9 domain was engineered to have a less stringent PAM requirement as compared to the parental version^50, 51^. ABE8e-editing of an HBG-175 target sequence was shown to be efficiently edit the expected adenine (A2) with potent HbF reactivation, as expected, and also produced concomitant bystander edits that either did not impact HbF expression (A3), or slightly decreased it (A1) (J.S.Y, M.J.W, T.M, pseronal communication). Delivery of HBG ABE8e mRNA and targeting gRNA by electroporation to human CD34^+^ cells resulted in the predicted A>G conversion at 3 different adenines within the target site, with efficiencies averaging 8% at A1, 40% at A2 and 18% at A3 (Fig6a-c). Similarly, to what we described earlier for CD33 editing, no indels or other types of changes were detected at this site (Fig. 6b). When used in the context of multiplex editing together with the CD33 ABE8e strategy described above, we noted a small drop in efficiency that did not reach statistical significance at all 3 adenines within the HBG target sequence (Fig. 6c). CD33 editing efficiency remained high in the context of multiplex editing, averaging 75% at A1 and 80% at A2 (Fig. 6d and Fig. 1c). Hematopoietic colony assays run from multiplex edited CD34^+^ cells revealed no difference in the number or type of colonies recovered relative to mock electroporated cells (Fig. 6e).

**Fig. 6.**
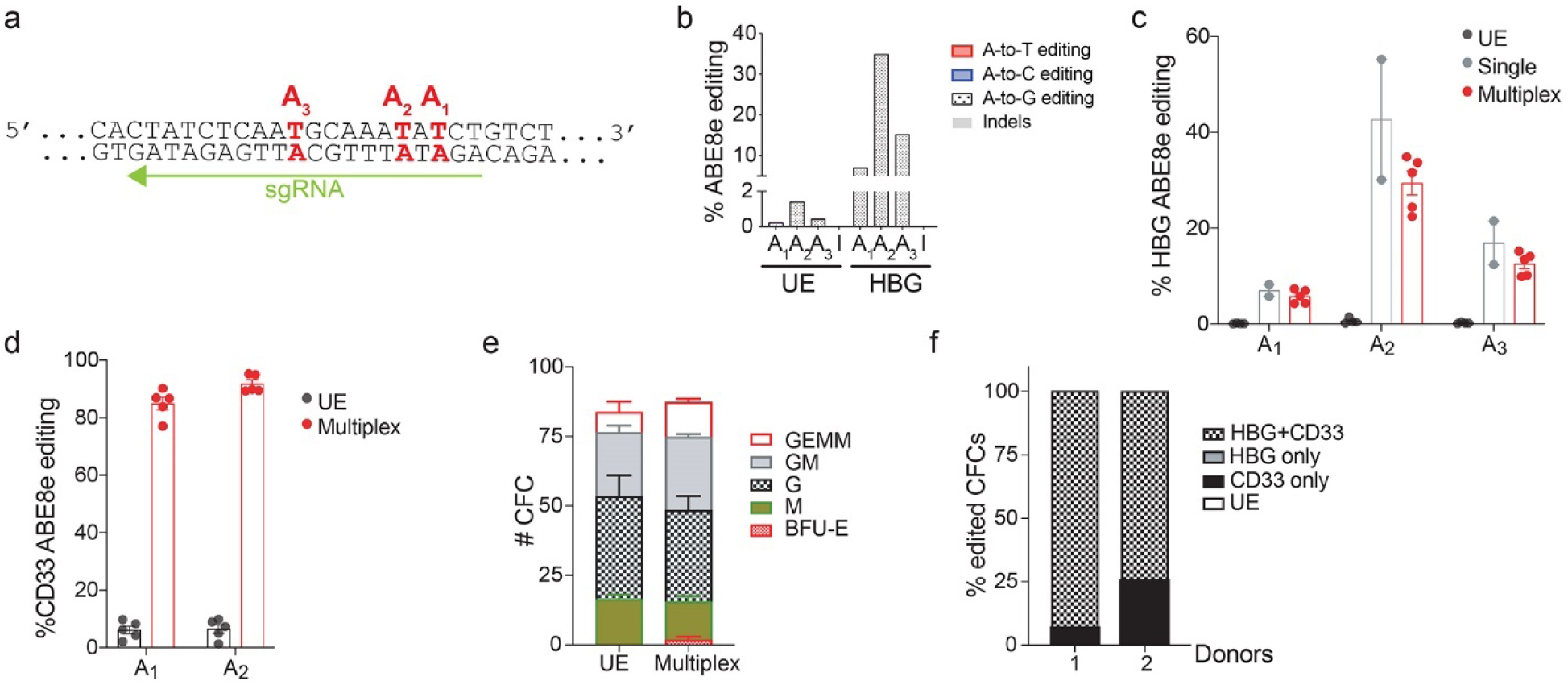
Multiplex CD33/HBG ABE8e editing of human CD34^+^ cells. a, Schematic of HBG -175 ABE8e target site. Adenines in editing window are marked in red and target site is shown with green arrow. b, HBG ABE8e editing efficiency measured in human mPB CD34^+^ HSPCs after treatment by mock electroporation (EP, WT) or by HBG ABE8e mRNA. Results are from one representative human donor. c, HBG editing efficiency in human mPB CD34^+^ HSPCs treated by mock EP (WT), by HBG ABE8e mRNA (single) or by CD33/HBG ABE8e mRNA EP (multiplex). Results are from 4 different human donors, 4 independent experiments. d, CD33 editing efficiency in human mPB CD34+ HSPCs treated by mock EP (WT) or by CD33/HBG ABE8e mRNA EP (multiplex). Results are from 5 different donors, 5 independent experiments. e, Colony-forming potential of mPB CD34^+^ cells treated by multiplex editing as described in c,d. Results are from 1 representative donor. f, Frequency of colony-forming cells displaying edits at each gene target, or at both gene targets in the same colony. Results are from 2 different human donors, 2 independent experiments.

Enrichment for therapeutically relevant HBG-edited cells using CD33-directed drugs will be feasible only if a substantial frequency of cells contains productive edits at both CD33 and HBG loci. To address this question, we assessed clonal editing outcomes in colonies derived from multiplex ABE8e edited CD34^+^ cells. Over 75% of colonies displayed edits at both the *HBG* and *CD33* genes (Fig. 6f). Of these multiplex edited colonies, the majority showed biallelic CD33 editing (100% and 78% in two different donors) and about half of the colonies display edits at two or more HBG genes (84.1% and 43.4%, Fig. S6). Taken together, these results indicate that ABE8e-mutiplex editing of mPB CD34^+^ HSPCs yields a large fraction of cells containing desired edits at both the HBG and CD33 gene targets, without compromising their multilineage differentiation capacity.

### Multiplex ABE8e-edited human CD34^+^ cells show normal hematopoietic reconstitution and GO resistance *in vivo*

We and others previous demonstrated that CRISPR/Cas9-editing of the *CD33* or *HBG* gene targets in CD34^+^ HSPCs does not compromise multilineage engraftment^1–3, 42^. We confirmed normal engraftment of CD33/HBG multiplex ABE8e-edited human CD34^+^ cells in the NSG mice, as well as multilineage differentiation in adult (Fig. 7) and neonate (Fig. S7) mice. In the adult model, human mPB CD34^+^ cell engraftment was comparable between the control group and the multiplex edited group at an average of 20% (Fig. 7a). As expected, CD33 expression in blood CD14^+^ monocytes were decreased to less than 10% in the edited group as compared to nearly 100% in the control group (Fig. 7a). Human lineage reconstitution was comparable between both groups in blood, spleen, and bone marrow at 16 weeks post-transplantation (Fig. 7a, b, d). When assessing bone marrow of mice receiving edited cells, CD33 was also almost completely absent from progenitors (Fig. 7d), including in the CD34^+^ subset (Fig. 7e). Importantly, HSC engraftment in bone marrow, as defined by CD34^+^CD38^low^, was not affecting by our editing approach (Fig. 7d). Together, these results demonstrate that our ABE8e-multiplex editing strategy does not affect CD34+ HSC multilineage engraftment in the mouse xenograft model.

**Fig. 7.**
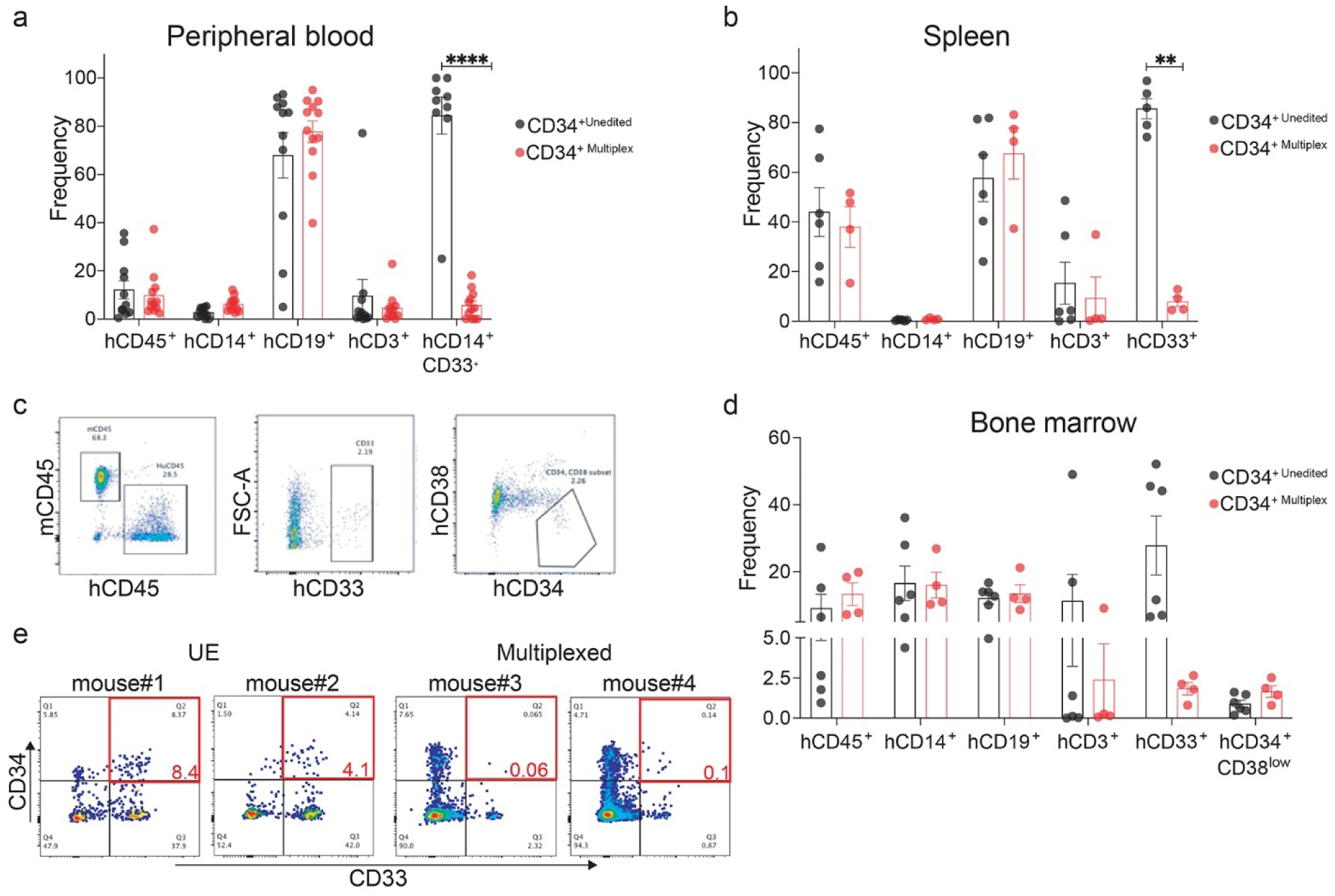
Engraftment of multiplex ABE8e-edited human mPB CD34+ cells in NSG mice. a,b, Measure of engraftment by percentage of human CD45^+^ cells and of hematopoietic repopulation by frequency of mature myeloid (CD14), lymphoid (CD19), and T cells (CD3) within the human CD45 population in peripheral blood (a) and spleen (b) at 14 to 16 weeks post-transplantation. CD33 expression within the CD14^+^ subset is also shown. For a, results are from 2 different human donors, 2 independent experiments, n=11 for WT and n=12 animals for multiplex editing. c, Representative flow plots showing gating strategy of human HSC (CD34^+^CD38^low^) in bone marrow at necropsy. d, Engraftment of multiplex edited vs. non-edited (WT) mPB CD34+ cells in bone marrow as measured by human CD45 expression and of hematopoietic repopulation by frequency of mature myeloid (CD14) and B and T lymphoid (CD19 and CD3) within the human CD45 population in peripheral blood at 14 to 16 weeks post-transplantation. CD33 expression in total bone marrow cells and HSC content (CD34^+^CD38^low)^ are also shown. Results are from 2 different human donors, 2 independent experiments, n=6 for WT and n=4 animals for multiplex editing. e, Representative flow plots of CD33 expression in bone marrow CD34^+^ cells at necropsy.

### Long-term persistence of multiplex ABE8e-edited CD34^+^CD90^+^ cells in NHPs

To evaluate the long-term safety and engraftment of multiplex ABE8e-edited HSPCs, we turned to the NHP model, which enables the investigation of multilineage engraftment, including myeloid differentiation, in an autologous setting. Furthermore, this model enables the tracking of hemoglobin production in peripheral blood, which is not possible in humanized mouse models where human erythroid cells do not fully mature^52^. We first verified that HSPC subsets, particularly the recently described HSC-enriched CD34^+^CD90^+^CD45RA^-^ population^53, 54^, could be efficiently edited with ABE8e in NHP. Bulk CD34^+^ cells were edited with CD33 ABE8e as described previously in human cells either using mRNA or RNP delivery, and subsequently sorted for HSPCs (CD90^+^CD45RA^−^), multipotent progenitor cells (CD90^−^CD45RA^−^), and lympho-myeloid progenitors (CD45RA^+^) (Fig. S8). CD33 editing was highest with mRNA delivery and was comparable for each sorted subpopulation, including for the HSC-enriched CD90^+^CD45RA^−^ subset, for both mRNA and RNP as delivery payload.

Having confirmed that the NHP HSC-enriched cell subset can efficiently be edited with ABE8e, we transplanted two rhesus macaques with multiplex edited cells. For both transplants, CD90^+^CD45RA^−^ cells were first FACS sorted from enriched bone marrow CD34^+^ cells (Fig. S8), which greatly minimized the number of cells to be manipulated. Consistent with our previous results employing CRISPR/Cas9 RNP-editing^42^ and despite upfront FACS-sorting, ABE8e-editing of a sorted CD90^+^ population did not affect CFC potential (Fig. S8). Editing efficiency in this cell population averaged 70 to 80% for the targeted A2 within CD33, and 8 to 45% for the three adenines at the HBG target site (Fig. 7a-b). Similarly, to human CD34^+^ cells (Fig. 6f), ∼70% of colonies derived from these cells contained edits at both target genes (Fig. 8c). Most colonies (83%) showed biallelic CD33 editing and about 40% of the colonies displayed at least two HBG edits (Fig. S7).

**Fig. 8.**
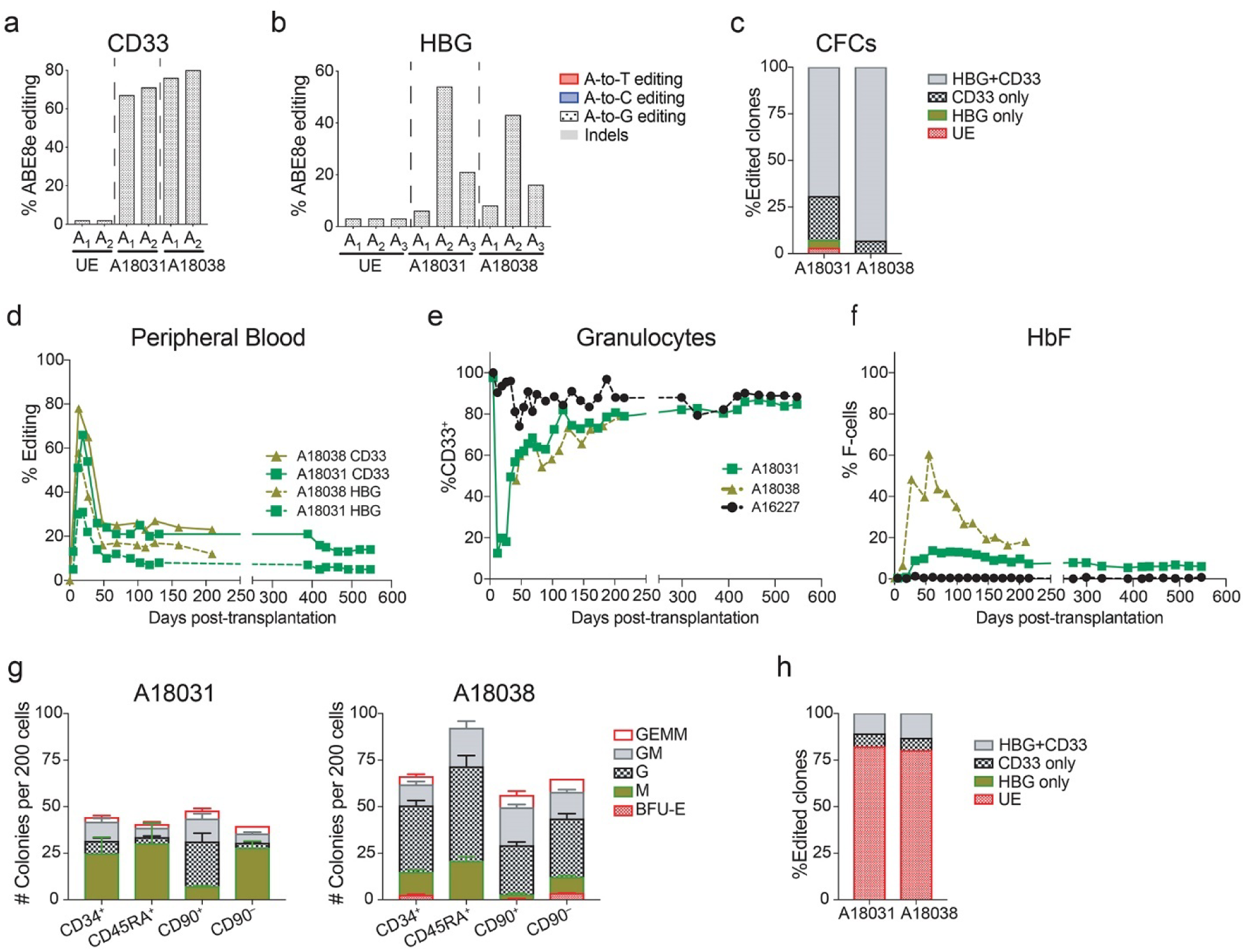
Long term engraftment of ABE8e multiplex edited CD90+ HSPCs in rhesus macaques. a,b, ABE8e editing efficiency at the CD33 and HBG targets measured in sorted rhesus CD90+ cells of each transplanted animal at 5 days post electroporation with ABE8e mRNA or in mock electroporated cells (WT). c, Frequency of colony-forming cells (CFCs) derived from cells edited in a and b displaying edits at each gene target, or at both gene targets in the same colony. n=57 for A18031 and n=45 for A18038. d, Tracking of CD33 and HBG editing in peripheral blood of both transplanted animals. For simplicity, only editing at A2 for CD33 or HBG is shown. e, CD33 expression in CD11b+CD14- peripheral blood granulocytes in both transplanted animals as compared to an untransplanted control animal (A16227). f, HbF reactivation as measured by flow cytometry staining for F-cells in peripheral blood of both transplanted animals as compared to the untransplanted control. g, CFCs derived from bone marrow aspirates taken from both transplanted animals at 8- or 6-months post-transplant for A18031 and A18038, respectively. h, Frequency of CFCs from g displaying edits at each gene target, or at both gene targets in the same colony. n=57 for A18031 and n=45 for A18038.

For animal A18031, the infused product consisted of total of 3.8×10^6^ edited CD90^+^CD45RA^−^ cells (454,000 cells per kg) combined with 10.6×10^6^ non-edited CD90^-^ cells (1.3×10^6^ cells per kg). For animal A18038, 7.56×10^6^ edited CD90^+^CD45RA^−^ cells (960,000 cells per kg) combined with 31×10^6^ non-edited CD90^-^ cells (3.9×10^6^ cells per kg). Both animals were conditioned by total body irradiation (TBI) and recovered without complications after transplantation. Blood counts including neutrophils, lymphocytes, platelets, and monocytes rapidly stabilized and remained within a normal range during posttransplant monitoring (Fig. S10). Hematopoietic recovery was also monitored by flow cytometry, which showed kinetics of multilineage reconstitution that are characteristic of a typical transplant experiment in this model^42, 53^ (Fig. S11). ABE8e editing at the CD33 and HBG gene targets in blood-nucleated cells stabilized after about 2 months post transplantation. At the last time point analyzed (18 months for A18031 and 7 months for A18038), CD33 editing at the relevant adenine (A2) reached 14% and 23%, respectively in each animal, while HBG editing at A2 was 5% and 12%, respectively (Fig. 8d). Resulting CD33 expression assessed by flow cytometry in blood granulocytes from each animal was lower post transplantation as compared to pre-transplant levels, or as compared to an un-transplanted animal control (Fig. 8e). In addition, the kinetics of CD33 expression post-transplant closely reflected the frequency of CD33 editing detected in blood nucleated cells in both animals. A similar correlation between HBG editing and frequency of blood F-cells was also noted, with levels that stabilized after the initial transient induction resulting from the transplantation procedure^55^ (Fig. 8f). At 6 months post-transplantation, 7% HBG editing resulted in 8% F-cells in animal A18031, and 16% HBG editing resulted in 16% F-cells in A18038 (Fig. 8d, f).

To verify that multiplex ABE8e-edited HSPCs are able to home back to the bone marrow after transplantation, BM samples were taken at 8- and 6-months post-transplantation in A18031 and A18038, respectively. Reconstitution of the stem cell compartment was assessed both by flow cytometry analysis and by functional readout using colony-forming assays. Immunophenotypic evaluation of BM samples demonstrated normal distribution of defined HSPC subsets that were not affected by our editing or transplant strategy (Fig. S12, S13). CFC assays demonstrated robust erythro-myeloid differentiation potential (Fig. 8g), similar to freshly isolated and nonmodified BM cells^42^. Single colonies derived from the CD90^+^ subset were also evaluated for editing at both CD33 and HBG as was done previously for the infusion product (Fig. 8c). About 20% of the colonies showed editing at one or both target genes, and importantly, 11% and 13% of the colonies showed editing at both targets in each animal, respectively (Fig. 8h). While these frequencies are lower than those detected in the infusion product, they confirm that multiplex ABE8e-edited HSPCs maintain multipotency and bone marrow homing capability. In summary, these results confirm that, in the autologous setting, CD90^+^-enriched HSPCs treated by ABE8e-multiplex editing are not compromised for long-term engraftment and differentiation.

### Multiplex ABE8e-edited human cells are enriched by GO treatment *in vitro*

Given that GO does not cross-react with rhesus CD33, we were unable to assess resistance and subsequent enrichment of multiplex ABE8e-edited cells to CD33 drugs in the NHP model. Instead, we used the ML1 human leukemic cell line that expresses CD33 and is highly sensitive to GO^3^. We demonstrated earlier that CD33 ABE8e-edited human CD34+ HSPCs are protected from GO treatment both in vitro (Fig. 2c) and in vivo (Fig. 3e, Fig. 7f-g). Following multiplex ABE8e-editing and GO treatment, we expect that cells containing single CD33 edits, and cells having both the CD33 and HBG edits to be selected for (Fig. 9a). Comparison of viable cell counts in mock electroporated cells (non-edited) with multiplex edited ML1 cells showed that edited cells were more resistant to the cytotoxic effect of the drug, and quickly proliferated after drug removal (Fig. 9b). Measurement of editing frequencies before and after GO treatment showed an increase of over 2-fold at the CD33 target site, and importantly, a 2-fold increase also at the HBG target (Fig. 9c). Together, these results confirm that treatment of multiplex edited cells with GO has the potential to increase levels of the HBG therapeutic edit. Future experiments will focus on evaluating this enrichment strategy in vivo.

**Fig. 9.**
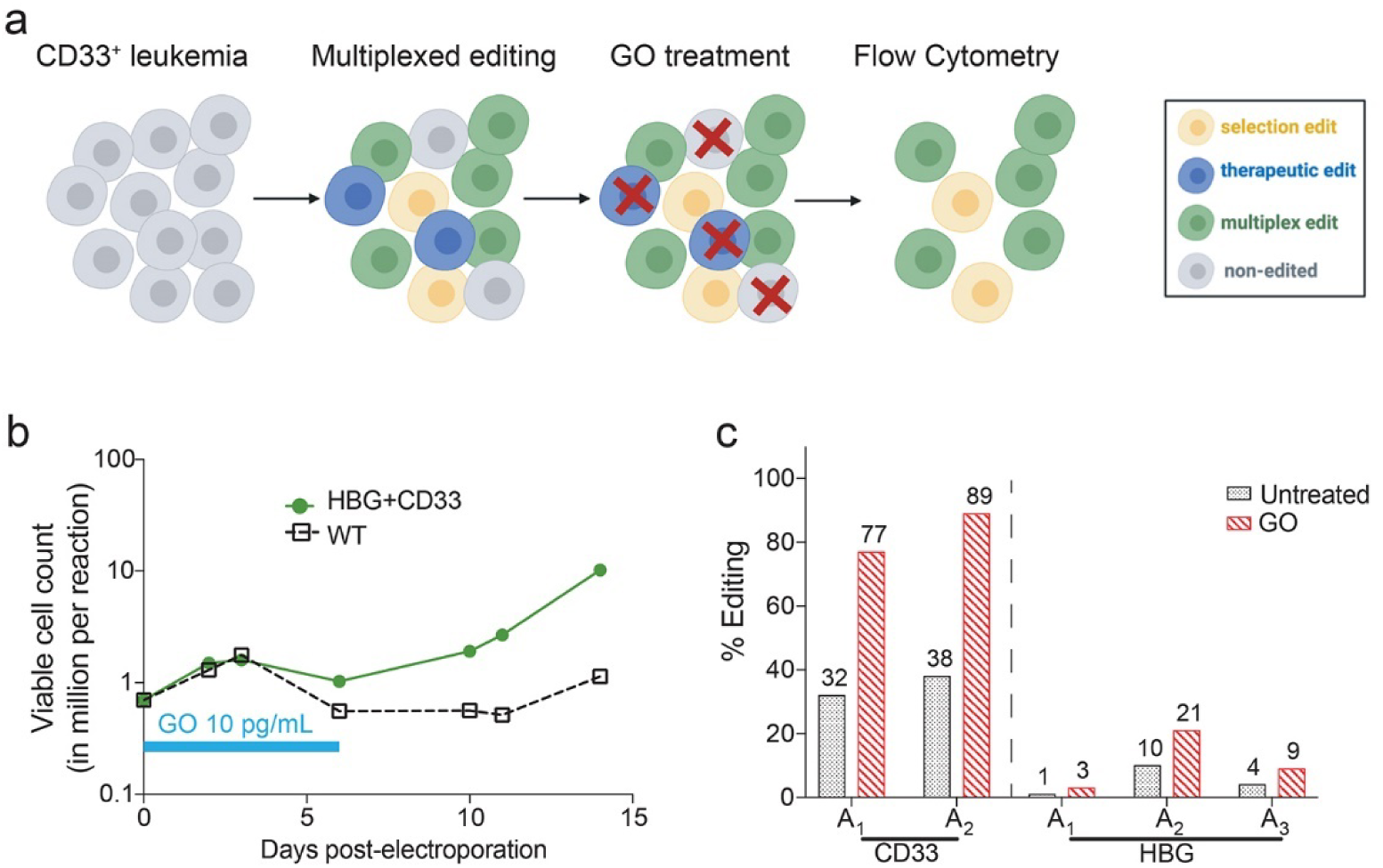
In *vitro* enrichment of multiplex ABE8e edited human ML1 cells. a, Schematic illustrating the principle of enrichment of multiplex edited cells after GO treatment. b, ML1 cells were mock electroporated (WT) or multiplex edited by ABE8e mRNA electroporation (HBG+CD33). At 5 days post-electroporation, cells were treated for 6 consecutive days with 10pg/mL GO. Viable count was determined by trypan blue staining. Results are from 1 independent experiment. c, Editing efficiency measured at the CD33 (left) or HBG (right) gene targets in multiplex edited ML1 cells treated or not with GO. Editing was measured at 8 days post GO removal (14 days post electroporation).

## Discussion

Here we use the adenine base editor ABE8e to disrupt the CD33 exon 2 splicing site as a strategy to protect cells from established CD33 immunotherapies targeting an epitope located in that exon. The safety of this approach was demonstrated by normal hematopoietic function of edited cells and by the lack of relevant off-target activity in engrafted HPSC. Our findings also showed resistance of CD33 ABE8e-edited HSPCs to GO both *in vitro* and *in vivo*, thus validating the concept of edited HSPCs as transplantation product that would be resistant to targeted therapies, using CD33 as a paradigm. To expand the applicability of this approach to other genetic diseases, particularly hemoglobinopathies, we next evaluated a multiplex CD33/HBG ABE8e editing strategy to determine if both edits could simultaneously be enriched following treatment with CD33-directed drugs. Since ABE edits DNA without introducing substantial DSBs, chromosomal translocations and toxicities associated with many DSBs are minimized, making ABE a particularly well-suited platform for multiplex genome editing. Multiplex CD33/HBG ABE8e edited HSPCs maintained normal and persisting multilineage engraftment in mouse and NHP models. In the latter model, we documented loss of CD33^FL^ surface expression in differentiated myeloid cells and elevated peripheral blood HbF production in two transplanted rhesus macaques for over a year. Given the lack of cross-reactive drugs with macaque CD33, we were unable to assess the proposed CD33 based enrichment strategy in the transplanted animals. Instead, we relied on a leukemic cell line to demonstrate *in vitro* that both CD33 and HBG edits were increased following treatment of the cells with GO.

The CD33 editing approach described in this report induces exon 2 skipping with few to no indels detected. By mimicking a natural SNP with no associated detrimental consequences, we generated HSPCs expressing an antigen variant lacking the exon containing the epitope recognized by the targeted therapy with no associated detrimental outcomes. CD33^Δ2^-edited cells exhibit normal function and differentiation but are insensitive to treatment with a CD33 antibody drug conjugate. Using a C-base editor to replicate the natural C>T SNP led to substantially increased indel outcomes (∼13%) and lower editing efficiency at the targeted nucleotides relative to an A-base editor that introduced a new mutation to destroy the same splice site. We expect that the high frequency of indels is caused by simultaneous deamination of 2 or more of the 4 C nucleotides in the editing window, recruiting uracil *N*-glycosylase that excises these modified nucleotides and results in double stranded DNA breaks together with the Cas9-induced nick^56^. Conversely, A-base editing achieved nearly 95% conversion of the targeted nucleotide within the splice acceptor site in HSPCs with no detectable indels. We propose to leverage the potential of allogeneic therapy for myeloid disorder by combining gene edited allo-HSCT with ADC and/or CAR immune cells.

We assessed off-target editing resulting from ABE8e treatment in HSPCs maintained in culture and in cells that engrafted in mice. Although some off-target editing was observed, there was no alteration of hematopoietic multipotency or myeloid function, which is promising for the clinical viability of this approach. Future research should focus on minimizing off-target editing. This could be achieved in part by reducing the concentration of editing reagents, since the selectable marker conferred by the CD33 edit could enrich edited cells from a population with lower overall editing.

Our group and others have shown that transplantation of CRISPR/Cas9 HBG-edited HSPCs stably increases HbF production while maintaining normal long-term hematopoietic reconstitution in vivo, including in the nonhuman primate (NHP) autologous model^42^. Similar CRISPR/Cas9-based approaches aimed at HbF reactivation are currently tested clinically^57^ for sickle cell anemia and beta-thalassemia^58^. Despite offering great promises, these approaches are limited by the use of myeloablative conditioning with busulfan for stable engraftment of gene-edited HSPCs. This use of high-intensity conditioning poses considerable risks such as infertility, organ toxicities and late-toxic effects, secondary malignancies, and prolonged myelosuppression with neutropenia. These limitations to HSC genome-editing based therapy could be alleviated using lower-intensity or antibody-based conditioning regimens. Given that these forms of conditioning may not be as efficient at removing non-edited endogenous cells, they are likely to result in lower levels of gene-modified cells, especially in diseases where edited HSPCs do not have a natural survival advantage such as hemoglobinopathies, an *in vivo* selection strategy is required to reach editing levels associated with therapeutic benefits. Given the resistance of CD33-edited cells to validated immunotherapies, our results showed *in vitro* that the secondary HBG edit is increased 2-fold. Further work is needed to demonstrate the feasibility of this approach *in vivo*, and whether selection will occur at the stem cell level or in differentiated myeloid cells expressing greater levels of CD33. The NHP model is ideally suited to evaluate selection strategy given the long-term persistence of modified cells but drugs that are cross-reactive with macaque CD33 are yet to be developed.

We propose an efficient editing strategy to create an HSC population that is both therapeutically altered and resistant to targeted CD33 therapy. Inspired by natural polymorphisms leading to exon skipping in CD33, we generated base editors that can lead to skipping of exon 2 that is targeted by anti-CD33 therapy. We propose that using genome editors to disrupt exons or otherwise modify residues involved in drug binding can be used to generate healthy cell populations resistant to many other cancer therapeutics. Adapting these methods to generate other resistance mutations, with or without multiplexed therapeutic edits, could improve targeted cancer therapies and treatments for genetic blood disorders.

## Material & Methods

### Base editor protein and mRNA production and purification

For protein expression, ABE8e and BE4max were codon optimized for bacterial expression and cloned into the protein expression plasmid pD881-SR (Atum, Cat. No. FPB-27E-269). The expression plasmid was transformed into BL21 Star DE3 competent cells (ThermoFisher, Cat. No. C601003). Colonies were picked for overnight growth in TB+25ug/mL kanamycin at 37 °C. The next day, 2L of pre-warmed TB were inoculated with overnight culture at a starting OD600 of 0.05. Cells were shaken at 37 °C for about 2.5 hours until the OD was ∼1.5. Cultures were cold shocked in an ice-water slurry for 1 hour, following which D-rhamnose was added to a final concentration of 0.8%. Cultures were then incubated at 18 °C with shaking for 24 hours to induce protein expression. Following induction, cells were pelleted and flash-frozen in liquid nitrogen and stored at -80 degrees. The next day, cells were resuspended in 30mL cold lysis buffer (1M NaCl, 100mM Tris-HCl pH 7.0, 5mM TCEP, 20% glycerol, with 5 tablets of cOmplete, EDTA-free protease inhibitor cocktail tablets (Millipore Sigma, Cat. No. 4693132001). Cells were passed 3 times through a homogenizer (Avestin Emulsiflex-C3) at ∼18000 psi to lyse. Cell debris was pelleted for 20 minutes using a 20,000xg centrifugation at 4 °C. Supernatant was collected and spiked with 40mM imidazole, followed by a 1-hour incubation at 4 °C with 1mL of Ni-NTA resin slurry (G Bioscience Cat. No. 786-940, prewashed once with lysis buffer). Protein-bound resin was washed twice with 12 mL of lysis buffer in a gravity column at 4 °C. Protein was eluted in 3mL of elution buffer (300 mM imidazole, 500 mM NaCl, 100 mM Tris-HCl pH 7.0, 5 mM TCEP, 20% glycerol). Eluted protein was diluted in 40 mL of low-salt buffer (100 mM Tris-HCl, pH 7.0, 5 mM TCEP, 20% glycerol) just before loading into a 50 mL Akta Superloop for ion exchange purification on the Akta Pure25 FPLC. Ion exchange chromatography was conducted on a 5 mL GE Healthcare HiTrap SP HP pre-packed column (Cat. No. 17115201). After washing the column with low salt buffer, the diluted protein was flowed through the column to bind. The column was then washed in 15mL of low salt buffer before being subjected to an increasing gradient to a maximum of 80% high salt buffer (1M NaCl, 100 mM Tris-HCl, pH 7.0, 5 mM TCEP, 20% glycerol) over the course of 50 mL using a flow rate of 5 mL per minute. 1 mL fractions were collected during this ramp to high salt buffer. Peaks were assessed by SDS-PAGE to identify fractions containing the desired protein, which were concentrated first using an Amicon Ultra 15 mL centrifugal filter (100 kDa cutoff, Cat. No. UFC910024), followed by a 0.5 mL 100 kDa cutoff Pierce concentrator (Cat. No. 88503). Concentrated protein was quantified using a BCA assay (ThermoFisher, Cat. No. 23227).

For mRNA production, ABE8e, ABE8e-NG and BE4max were ordered as custom products from Trilink Biotechnologies. mRNAs were transcribed in vitro from PCR product using full substitution of N1-methylpseudouridine for uridine and was capped co-transcriptionally using the CleanCap AG analogue (Trilink Biotechnologies), resulting in a 5’ Cap 1 structure. The in vitro transcription reaction was performed as previously described.^15, 59^. Mammalian -optimized UTR sequences (Trilink) and a 120-base poly-A tail were included in the transcribed PCR product.

### Culture and editing of CD34^+^ HSPCs

CD34^+^ Cord Blood cells were obtained from de-identified healthy adult donors (Stemexpress, or Lonza). For mobilized peripheral blood CD34^+^ cells, primary human CD34^+^ cells were purchased from the Co-operative Center for Excellence in Hematology (CCEH) at the Fred Hutchinson Cancer Center. Collections were performed according to the Declaration of Helsinki and were approved by a local Ethics Committee/Institutional Review Board of the Fred Hutchinson Cancer Center. All healthy adult donors were mobilized with granulocyte colony-stimulating factor (G-CSF). Human CD34^+^ cells were enriched as previously described on a CliniMACS Prodigy according to the manufacturer’s instructions (Miltenyi Biotec). Nonhuman primate CD34^+^ cells were harvested, enriched, and cultured as previously described^42, 53^. Briefly, before enrichment of NHP CD34+ cells, red cells were lysed in ammonium chloride lysis buffer, and WBCs were incubated for 20 minutes with the 12.8 immunoglobulin-M anti-CD34 antibody, then washed and incubated for another 20 minutes with magnetic-activated cell-sorting anti-immunoglobulin-M microbeads (Miltenyi Biotech). Cells were cultured and expanded in StemSpan SFEM II (StemCell technologies) containing 1% Penicillin Streptomycin, 100 ng/mL TPO, 100 ng/mL SCF, 100 ng/mL IL6 and 100 ng/mL FLT3L at 37°C, 85% relative humidity, and 5% CO2 (human cytokines from Biolegend or PeproTech). Cord blood CD34^+^ cells were additionally cultured with UM171 0.35 nM (Xcessbio). In colony-forming cell assays, 1000-1200 sorted cells were seeded into 3.5 ml ColonyGEL 1402 (ReachBio).

Chemically modified sgRNA were purchased from Synthego. For base editor protein, RNP were formed by mixing 5 to 9 μgr base editor protein with 1.5 μgr of sgRNA in 20 μL of P3 buffer (Lonza, Amaxa P3 Primary Cell 4D-Nucleofector Kit) and incubated for 10 mins. Then cells were electroporated at 50 million cells per mL. For multiplex editing using base editor mRNA, electroporation reactions were generally conducted using 3 μg mRNA combined with 100 pmol of each sgRNA for 1 million cells using the ECM 380 Square Wave Electroporation system (Harvard Apparatus).

### *In vitro* phagocytosis assay

CD34^+ Unedited^ cells or CD33^Δ2^-edited CD34^+^cells were differentiated into monocytes for 14 days, with StemSpan SFEM II containing 1% Penicillin Streptomycin and the StemSpan Myeloid Expansion Supplement II (StemCell Technologies). At day 15, differentiated monocytes were preincubated for 30 mins with PBS or 25 μμM Cytochalasin D and then incubated with pHrodo red E coli bioparticles (Thermofisher) with or without 25 μM cytochalasin D for 1 h 30 mins. Cells were then transferred to FACS tubes, washed with PBS 1% FBS and incubated 10 mins at RT with Human TruStain FcX and True-Stain Monocyte blocker (Biolegend). Antibody mix containing CD14-APCFire750, CD13-BV786, CD34-BV421, CD33-APC (Biolegend) and ebioscience fixable viability dye e450 (Thermofisher) was then added directly. After 25 mins of incubation, cells were washed and phagocytosis assessed by FACS.

### *In vitro* cytotoxicity assay

CD34^+Unedited^, CD33^Δ2^-edited CD34^+^, CD34^+^ cells carrying heterozygous or homozygous rs12459419 SNP or MOLM14 cells were incubated in triplicate with or without GO for 24 or 48 hours. Cells were then washed and stained with viability dye 7AAD (Biolegend) or Sytox blue (Thermofisher) and immediately analyzed by FACS. CD3/CD33 BiAb-induced cytotoxicity was determined as previously described^3^. Briefly, AML cells were incubated at 37°C (in 5% CO_2_ and air) in 96-well round-bottom plates (BD Biosciences) at 5 to 10 × 10^3^ cells per well containing various concentrations of the Bispecific T cell engager (generated from published sequences using methods similar to those reported previously^60^, as well as T cells at different effector to target (E:T) cell ratios. After 48 hours, cell numbers and drug-induced cytotoxicity, using 4’,6-diamidino-2-phenylindole (DAPI) to detect non-viable cells, were determined by FACS (BD Biosciences).

### CIRCLE-seq off target editing analysis

To identify base editing off target sites, genomic DNA was extracted from CD34^+^ cells with the QIAgen Gentra PureGene kit. CIRCLE-seq was performed as previously described^39^. Briefly, purified genomic DNA was sheared with a Covaris S2 instrument to an average length of 300 bp. The fragmented DNA was end repaired, A-tailed and ligated to an uracil-containing stem-loop adaptor, using KAPA HTP Library Preparation Kit, PCR Free (KAPA Biosystems). Adaptor ligated DNA was treated with Lambda Exonuclease (NEB) and E. coli Exonuclease I (NEB) and then with USER enzyme (NEB) and T4 polynucleotide kinase (NEB). Intramolecular circularization of the DNA was performed with T4 DNA ligase (NEB) and residual linear DNA was degraded by Plasmid-Safe ATP-dependent DNase (Lucigen). *In vitro* cleavage reactions were performed with 250 ng of Plasmid-Safe-treated circularized DNA, 90 nM of Cas9 nuclease (NEB), Cas9 nuclease buffer (NEB) and 90 nM of synthetically modified sgRNA (Synthego) in 100 μL volume. Cleaved products were A-tailed, ligated with a hairpin adaptor (NEB), treated with USER enzyme (NEB) and amplified by PCR with barcoded universal primers NEBNext Multiplex Oligos for Illumina (NEB), using Kapa HiFi Polymerase (KAPA Biosystems). Libraries were sequenced with 150 bp paired-end reads on an Illumina MiSeq instrument. CIRCLE-seq data analyses were performed using open-source CIRCLE-seq analysis software (https://github.com/tsailabSJ/circleseq) using default parameters. The human genome GRCh37 was used for alignment.

### Quantification of base-editing efficiency at on-target and off-target sites with High throughput sequencing (HTS) using the Illumina MiSeq

To assess the on-target and off-target editing, we sequenced the CD33 locus and the 19 top off-target sites nominated by CIRCLE-seq. Genomic DNA from CD34^+^ cells or Bone Marrow of transplanted mice was extracted with the QuickExtract™ DNA Extraction Solution (Lucigen). For CD33 and each off-target site, we designed primers to generate a 250-300bp product including the aligned off-target binding site for the guide RNA, and appended adapters (in italic) for Illumina sequencing. To investigate ABE8e mediated editing, the CD33 locus was amplified from genomic DNA using the following primers: Forward 5’-*ACACTCTTTCCCTACACGACGCTCTTCCGATCTNNNN*CCTCCACTCCCTTCCTCTTT-3’and the Reverse 5’-*TGGAGTTCAGACGTGTGCTCTTCCGATC*TCTTCCCGGAACCAGTAACCA-3’. To investigate BE4max/sgCBE_1_ mediated editing, the CD33 locus was amplified with the Forward 5’-*ACACTCTTTCCCTACACGACGCTCTTCCGATCTNNNN*AGACATGCCGCTGCTGCTA-3’ and the Reverse 5’-*TGGAGTTCAGACGTGTGCTCTTCCGATC*CCGGAACCAGTAACCATGAAC-3’. Following a secondary PCR to barcode each sample, products were pooled and sequenced using a 300 cycle v2 Illumina MiSeq kit. To analyze editing outcomes in resulting fastq files, we used CRISPResso2^61^ to align each read to the reference amplicon and quantify indels or base changes.

In the multiplex editing experiments, editing was quantitated by next generation sequencing (NGS) or by EditR as follow. Genomic DNA was extracted using QIAamp DNA micro kit (Qiagen, Germantown, MD, U.S.), and processed by PCR amplification using the primers described in Table S1. In NGS, libraries were prepared using Illumina barcoded, 2×150 base-pair (bp), pair-end and run on the Miseq platform (Illumina, San Diego, CA, USA). In TIDE analysis, amplicons were sequenced by Sanger sequences and analyzed using the web-based EditR program^62^. The raw bioinformatic data and custom R code generated for the analysis of editing efficiency are available on request.

### Mouse transplantation experiments

Mice experiments were either conduced at the Columbia University animal facility under specific pathogen-free (single CD33 editing) or at the Fred Hutchinson Cancer Center in compliance with the approved Institutional Animal Care and Use Committee (IACUC) protocol #50980.

#### Columbia University

NOD.Cg-Prkdcscid Il2rgtm1Wjl Tg(CMV-IL3,CSF2,KITLG)1Eav/MloySzJ (NSGS) or NSG mice (The Jackson Laboratory) were conditioned with sublethal (1.2 Gy) total-body irradiation (TBI). Human Cord Blood stem cells (1×10^6^per mouse) were injected intravenously into the mice within 8-24 hours post-TBI. Engraftment and repopulation of the hematopoietic system over time, was assessed by analysis of peripheral blood, whole bone marrow and spleen using the consequent antibodies from Biolegend or BD Biosciences: Ter119-PeCy5, Ly5/H2kD-BV711, hCD45-BV510, hCD3-Pacific Blue, hCD123-BV605, hCD33-APC, hCD14-APC/Cy7, hCD10-BUV395, hCD19-BV650. Dead cells were excluded using Propidium Iodide. For *in vivo* GO treatment, engraftment and hematopoietic repopulation was assessed in bone marrow (aspiration) and peripheral blood. Mice were then injected intravenously with GO (resuspended in water) at the indicated concentration. On week after treatment, bone marrow and peripheral blood were analyzed for the presence of myeloid cells by FACS. All data were acquired with the Bio-Rad ZE5 flow cytometry analyzer and data analysis were performed using FlowJo 10.8.1. All *in vivo* experiments were performed under protocols approved by the institutional Animal Care and Use Committee at Columbia University.

#### Fred Hutchinson Cancer Center

NSG (non-obese diabetic [NOD].Cg-PrkdcscidIl2rgtm1Wjl/SzJ) mice were irradiated at 275 cGy. 4 hours later, mice were intravenously or intrahepatically injected with 1×10^6^ (adult model) or 0.3×10^6^ (neonate model) CD34+ cells, respectively, resuspended in 1X PBS with 1% heparin (APP Pharmaceuticals, East Schaumburg, IL, U.S.) to a total volume of 200 μl (adult) or 30 μl (neonate), preceded by sublethal irradiation of 275 cGy (adult) or of 150 cGy (neonate).

Beginning at 8 weeks post-injection, blood samples were collected every 2 to 4 weeks and analyzed by flow cytometry. At time of necropsy, animals were sacrificed, and tissues harvested and analyzed. Lineage and HSC assessment from blood or BM were performed by flow cytometry utilizing antibody panel described in Table S2, and run on the BD FACSCanto II or BD Symphony Flow Cytometer (BD Biosciences). Data were acquired using FACSDiva version 6.1.3 and newer (BD Biosciences). Data analysis was performed using FlowJo version 8 and higher (BD Biosciences). For BM CFCs assays, 7×10^4^/ml cells were plated in ColonyGEL^TM^ 1402 (ReachBio, Seattle, WA, U.S.) in triplicate from each mouse BM. Colonies were counted and picked for genomic analysis after 14 days.

### NHP transplant experiments

Autologous NHP transplants were conducted as described previous for CRISPR/Cas9-editing of the CD90^+^ subset^42^. Briefly, rhesus macaques were conditioned with myeloablative total body irradiation (TBI) of 1020 cGy from a 6 MV x-ray beam of a single-source linear accelerator located at the Fred Hutch South Lake Union Facility (Seattle, Washington, USA). The dose was administered at a rate of 7 cGy/min delivered as a midline tissue dose. After transplant, G-CSF was administered daily from the day of cell infusion until the animals began to show onset of neutrophil recovery. Supportive care, including antibiotics, electrolytes, fluids, and transfusions, was given as necessary, and blood counts were analyzed daily to monitor hematopoietic recovery. All experimental procedures performed were reviewed and approved by the Institutional Animal Care and Use Committee of the Fred Hutchinson Cancer Center (Fred Hutch) and University of Washington (UW; Protocol #3235-01). This study was carried out in strict accordance with the recommendations in the *Guide for the Care and Use of Laboratory Animals of the National Institutes of Health* (“The Guide”), and monkeys were randomly assigned to the study. Editing from blood and bone marrow was tracked in the transplanted animals as described in the section above, and HbF and multilineage reconstitution was monitored by flow cytometry as described previously^42, 55^ and using the antibodies listed in Table S2.

#### Nonhuman primate animal housing and care / ethics statement

Healthy juvenile pigtailed macaques and juvenile rhesus macaques were housed at the UW National Primate Research Center (WaNPRC) under conditions approved by the American Association for the Accreditation of Laboratory Animal Care. All experimental procedures performed were reviewed and approved by the Institutional Animal Care and Use Committee of Fred Hutch and UW (Protocol #3235-01). This study was carried out in strict accordance with the recommendations in the Guide for the Care and Use of Laboratory Animals of the National Institutes of Health (“The Guide”) and monkeys were randomly assigned to the study. This study included at least twice-daily observation by animal technicians for basic husbandry parameters (e.g., food intake, activity, stool consistency, and overall appearance), as well as daily observation by a veterinary technician and/or veterinarian. Animals were housed in cages approved by “The Guide” and in accordance with Animal Welfare Act regulations. Animals were fed twice daily and were fasted for up to 14 hours prior to sedation. Environmental enrichment included grouping in compound, large activity, or run-through connected cages, perches, toys, food treats, and foraging activities. If a clinical abnormality was noted by WaNPRC personnel, standard WaNPRC procedures were followed to notify the veterinary staff for evaluation and determination for admission as a clinical case. Animals were sedated by administration of ketamine HCl and/or telazol and supportive agents prior to all procedures. Following sedation, animals were monitored according to WaNPRC standard protocols. WaNPRC surgical support staff are trained and experienced in the administration of anesthetics and have monitoring equipment available to assist with electronic monitoring of heart rate, respiration, and blood oxygenation; audible alarms and LCD readouts; monitoring of blood pressure, temperature, etc. For minor procedures, the presence or absence of deep pain was tested by the toe-pinch reflex. The absence of response (leg flexion) to this test indicates adequate anesthesia for this procedure. Similar parameters were used in cases of general anesthesia, including the loss of palpebral reflexes (eye blink). Analgesics were provided as prescribed by the Clinical Veterinary staff for at least 48 hours after the procedures and could be extended at the discretion of the clinical veterinarian, based on clinical signs.

### Statistics

All numerical results are reported as mean ± SEM. All statistical analyses were performed with GraphPad Prism 9.3.1. Statistical significance of the difference between experimental groups were analyzed mainly with unpaired t-tests or one-way ANOVA, unless otherwise stated in the figure legend. Differences were considered statistically significant for p≤0.05 and denoted as: *p≤0.05; **p≤0.01; ***p≤0.001; ****p≤0.0001.

### Data availability

HTS sequencing files can be accessed on the NCBI Sequence Read Archive.

## Acknowledgments

Flow cytometry experiments were performed using instrumentation maintained by the Columbia Stem Cell Initiative Flow Cytometry core facility directed by Michael Kissner and by the Fred Hutchinson Cancer Center. We are grateful to Veronica Nelson, Erica Wilson, Kelvin Sze, Megan Brown, Sarah Herrin, Alan Ung, and Michelle Hoffman for outstanding support in our NHP studies. All primate work was completed at the WaNPRC, which is supported by U42 OD011123 and P51 OD010425 grants through the NIH Office of Research Infrastructure Programs. G.A.N. was supported by a Helen Hay Whitney Postdoctoral Fellowship and NIH K99 award HL163805. H-P.K. acknowledges support from NIH R01 HL136135. D.R.L. acknowledges support from NIH U01 AI142756, RM1 HG009490, R35 GM118062, and HHMI. R.B.W acknowledges support from NIH/NHLB R01 HL151765 and NIH/NCI R01 CA266556. M.J.W. acknowledges support from NIH P01 HL053749, R01 HL156647, and from ALSAC. We thank Mallory Llewellyn and Savannah Cook for excellent technical assistance. Biorender was used to create all the diagrams, cartoons and schematics shown along the manuscript, under the Columbia University or Fred Hutchinson Cancer Research Center academic licenses. We are grateful to Helen Crawford for help in preparing this manuscript and figures.

## Authors contributions

F.B, O.H, E.F, G.A.N, S.R, G.S.L. and S.K designed, performed experiments and analyzed data. A.M.A, R.W, M.J.W, J.S.Y, T.M provided conceptual assistance. F.B, O.H, S.M and H-P.K wrote the manuscript with input from all authors. S.M and H-P.K supervised the study.

## Supplemental Data

**Supplemental Fig. 1.**
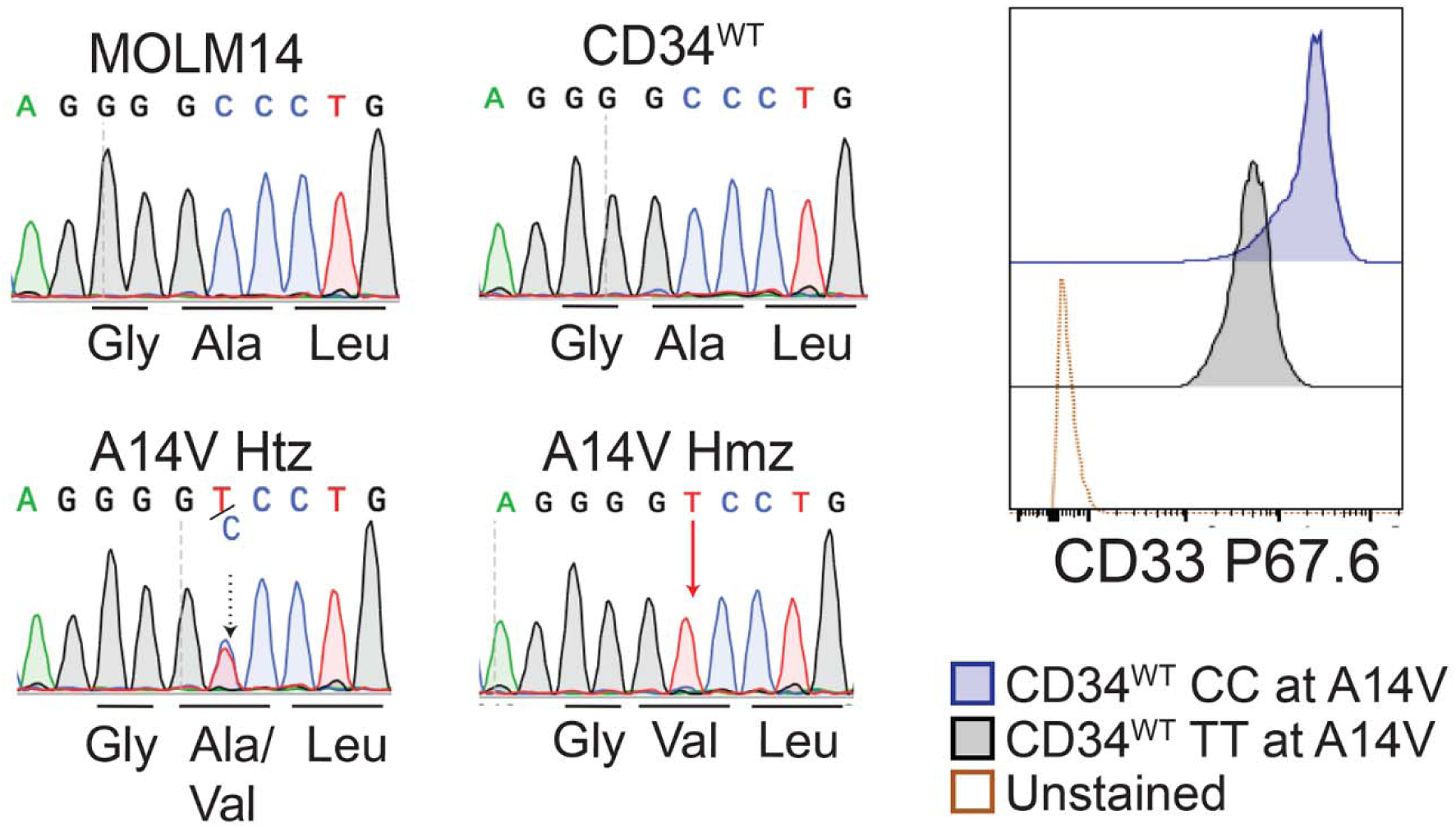
Sequencing profile of MOLM14 (AML immortalized cell line), CD34^+WT^ (unedited cells in this study) and two donors with heterozygous (CT) or homozygous (TT) rs12459419 A14V SNP. (C) FACS profile of two CD34^+^ donors homozygous CC or TT at A14V.

**Supplemental Fig. 2.**
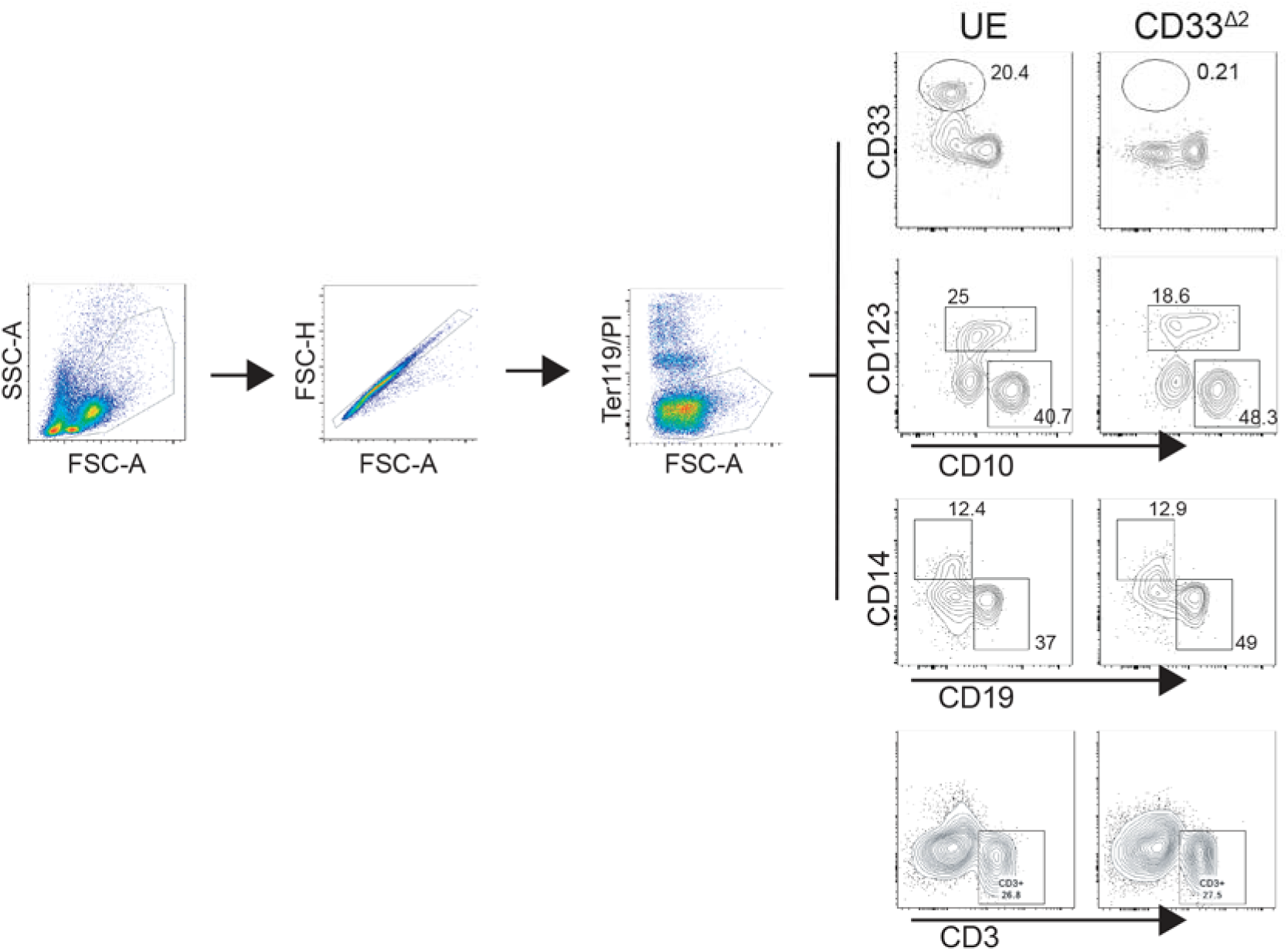
Gating strategy for in vivo assesment (UE, Unedited)

**Supplemental Fig. 3.**
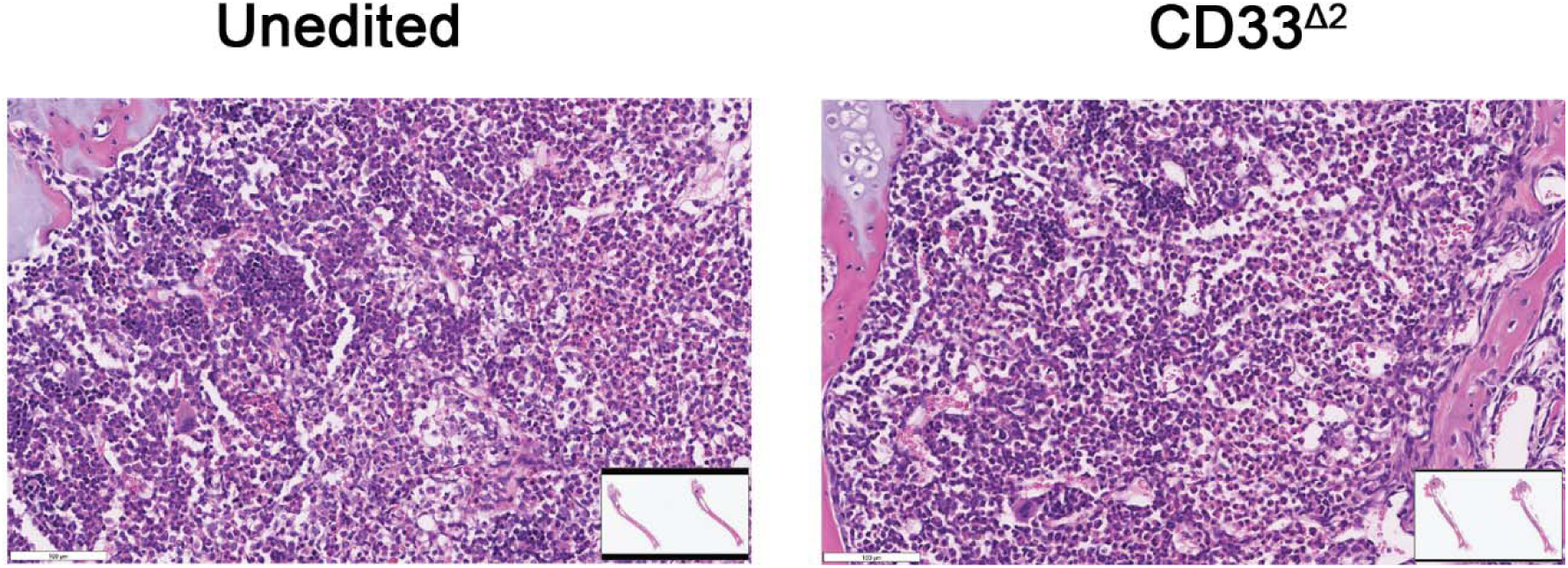
H&E staining of Bone Marrow of mice engrafted with CD34^+ unedited^ or CD34^+^CD33^Δ2^ cells, at 16 weeks post-transplantation.

**Supplemental Fig. 4.**
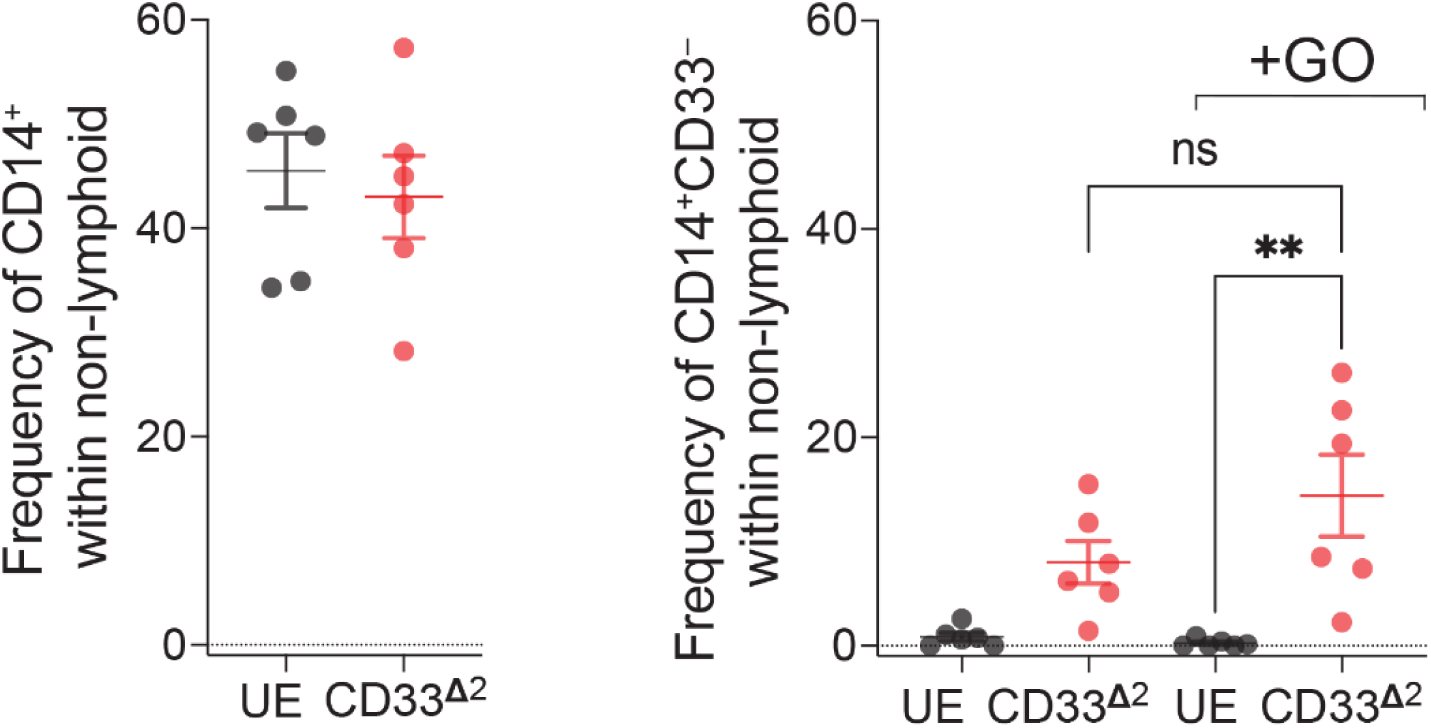
Left, frequency of CD14^+^ myeloid cells in BM of 12 weeks post-transplanted mice *Right*, frequency of CD14^+^CD33^+^ cells in the BM of 12 weeks post-transplanted mice before and one week after GO treatment (0.5 ug per mouse). UE, unedited cells. One way ANOVA.

**Supplemental Fig. 5.**
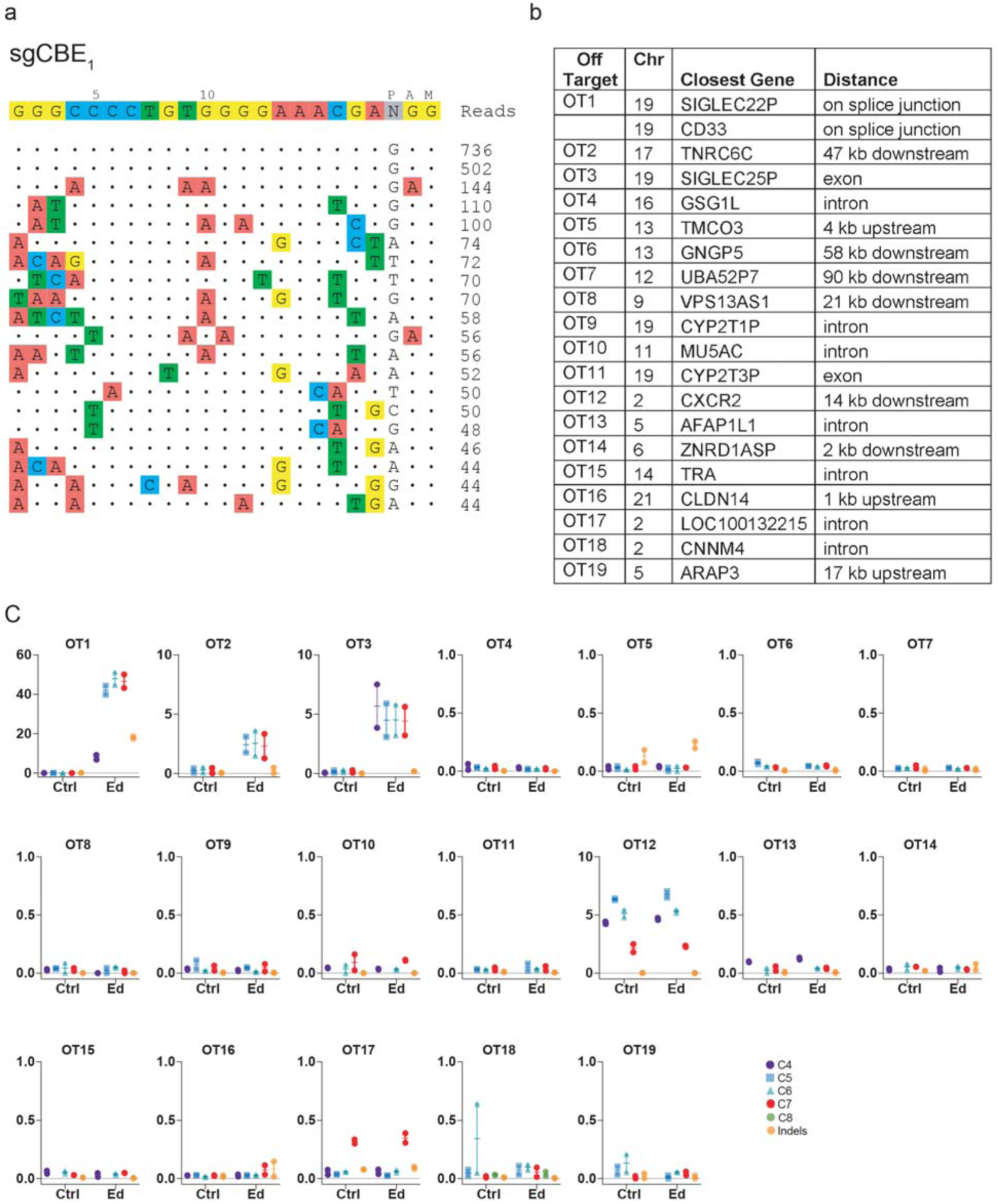
Off target associated with BE4max protein editing of CD33 exon 2 ESE in CD34^+^ HSPCs kept *in vitro.* **a,** Visualization of on target and top 20 off target sites detected by CIRCLE- seq. Read counts and alignement of each site to the sgCBE1 protospacer is shown **b,** Table summarizing the 19 top off target identified loci. **c,** Editing assessment of C-to-T editing at position C3,C4,C5,C6,C7 and indels at the 19 identified top off target loci in CD34^+^ HSPCs kept *in vitro* unedited (control, Ctrl) or edited (Ed) cells. Amplicons were sequenced by HTS and C-to-T editing was quantified. Data shown as mean ± SEM.

**Supplemental Fig. 6.**
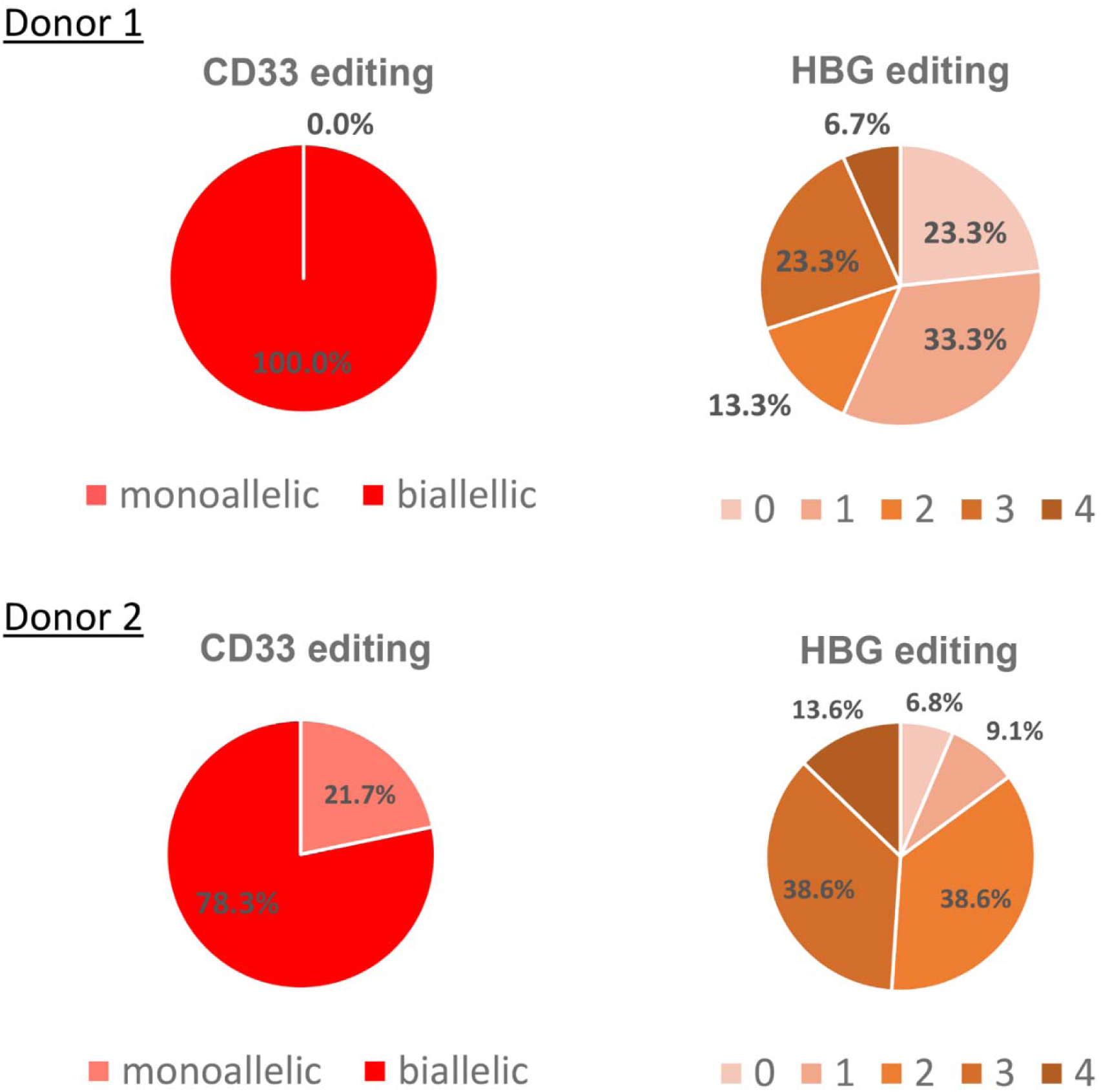
Detail of monoallelic vs. biallelic editing at the HBG and CD33 targets in single colonies derived from multiplex edited human CD34+ cells used in the mouse studies (Fig. 7). PCR amplicons for each target was produced from single colony lysates, sequenced by Sanger sequencing, and analyzed by EditR.

**Supplemental Fig. 7.**
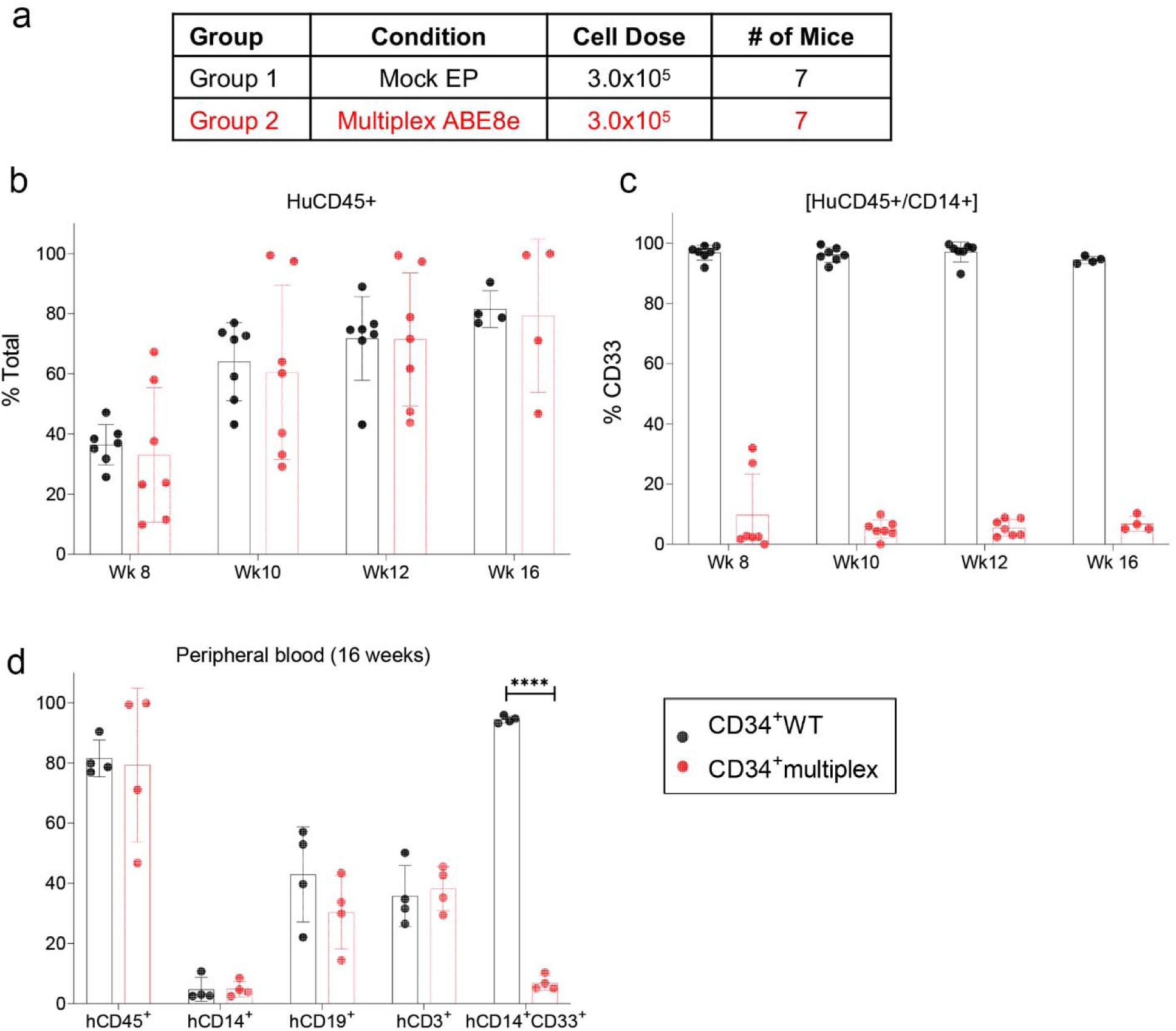
Multiplex ABE8e-edited CD34+ cells engraftment in the mouse neonate model. **a,** Details of experiment layout. **b,** Measure of engraftment by percentage of human CD45^+^ cells in peripheral blood. **c,** CD33 expression measured within the CD14^+^ subset from peripheral blood. **d,** Hematopoietic repopulation by frequency of mature myeloid (CD14), lymphoid (CD19), and T cells (CD3) within the human CD45 population in peripheral blood from animals of both experimental groups.

**Supplemental Fig. 8.**
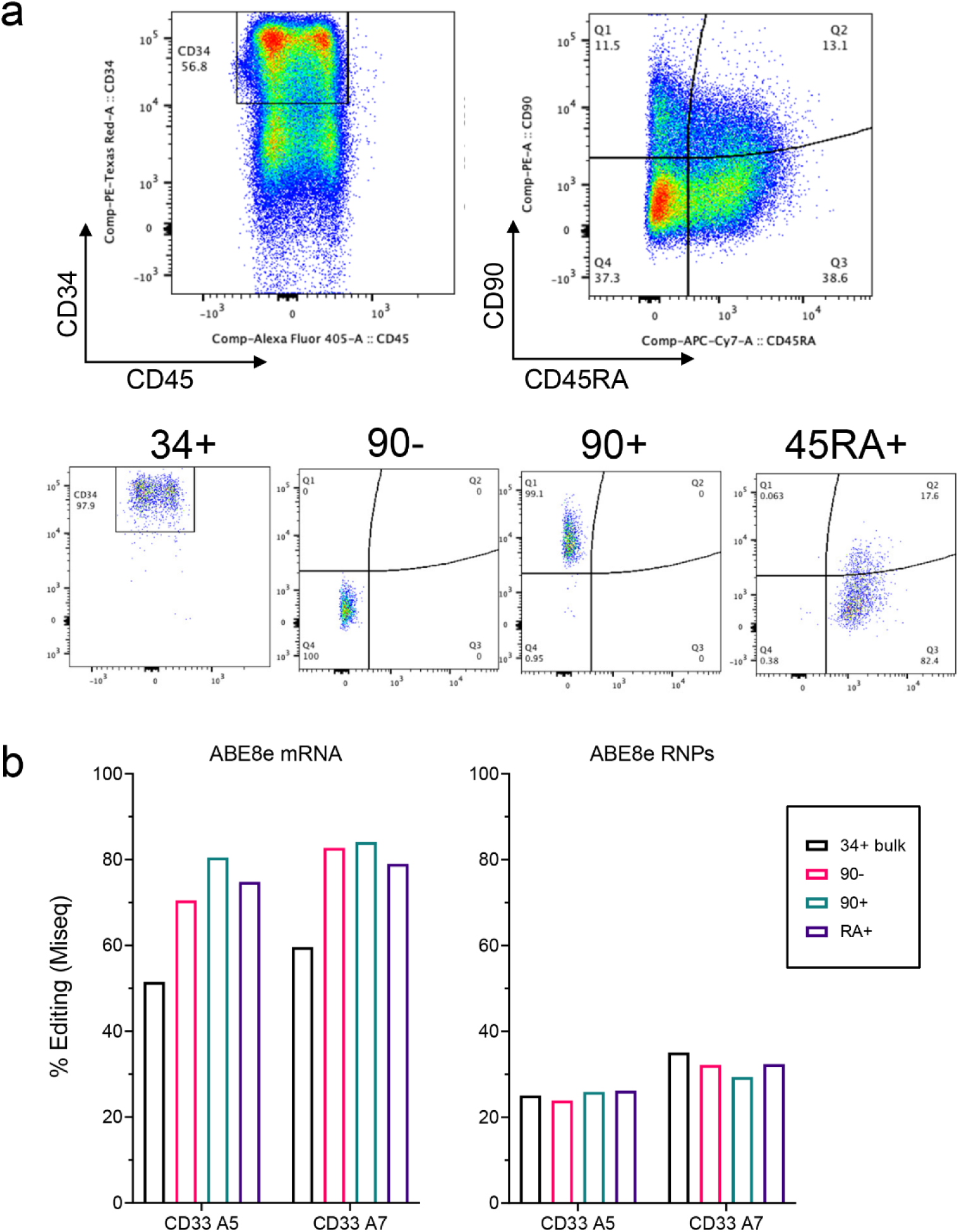
CD33 editing using ABE8e mRNA or protein in different HSPC subsets in NHP. **a,** Representative flow cytometry plots showing gating strategy for the different subset before (top) and after (bottom) sort. **b,** Editing efficiency measured by NGS in sorted subsets after ABE8e mRNA (left) or protein (right) treatment.

**Supplemental Fig. 9.**
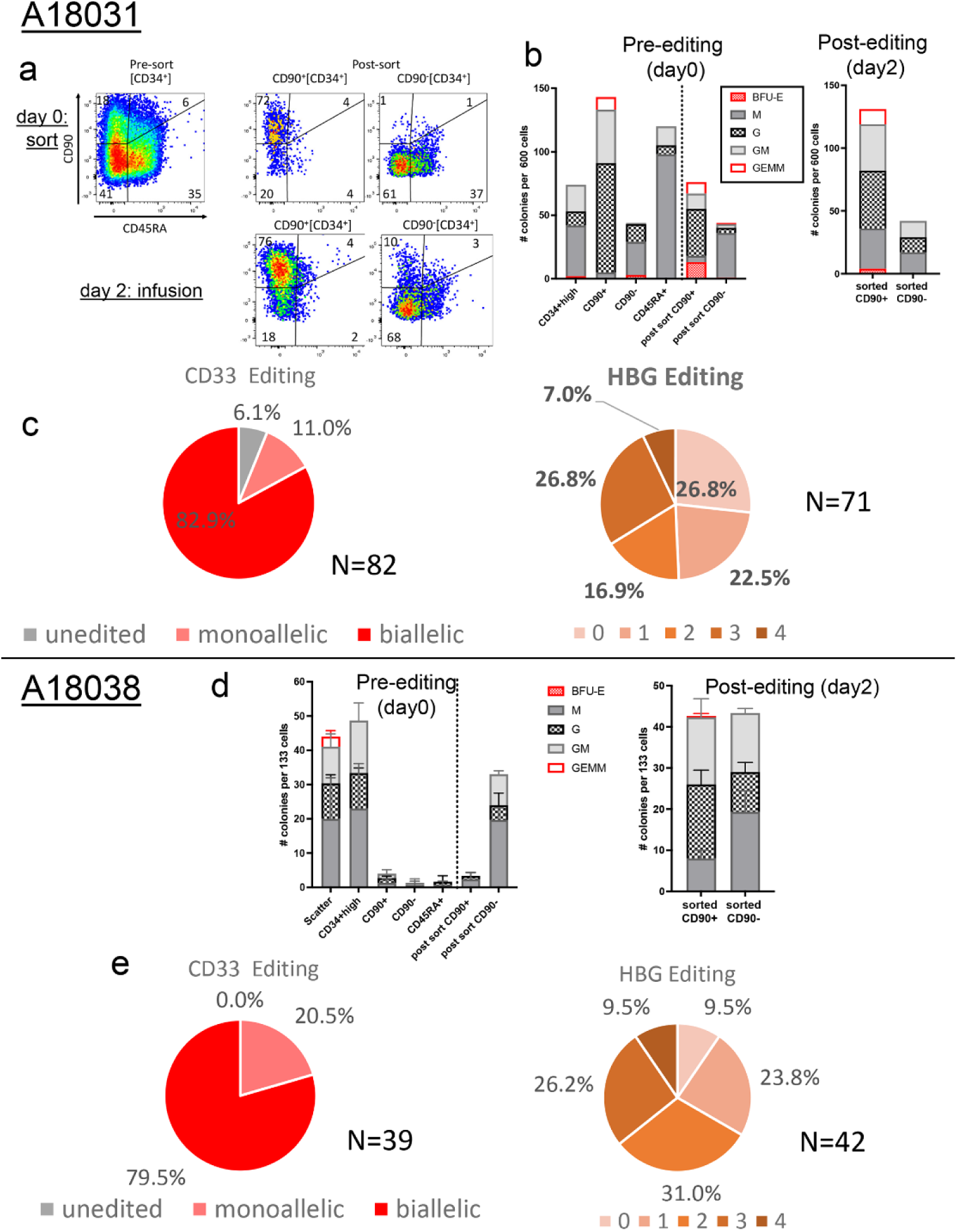
Validation of NHP infusion products by flow cytometry and colony-forming assays for both transplanted animals and assessment of monoallelic vs. biallelic editing in single colonies. **a,** Phenotypic characterization of infusion product by flow cytometry plots at the time of HSC purification (day 0) or at the time of infusion (day 2). **b,** Colony-forming assay of the different subsets shown in (a) at day 0 or day 2. **c,** Detail of monoallelic vs. biallelic editing at the HBG and CD33 targets in colonies derived from multiplex edited NHP HSPCs. PCR amplicons for each target was produced from single colony lysates, sequenced by Sanger sequencing, and analyzed by EditR.

**Supplemental Fig. 10.**
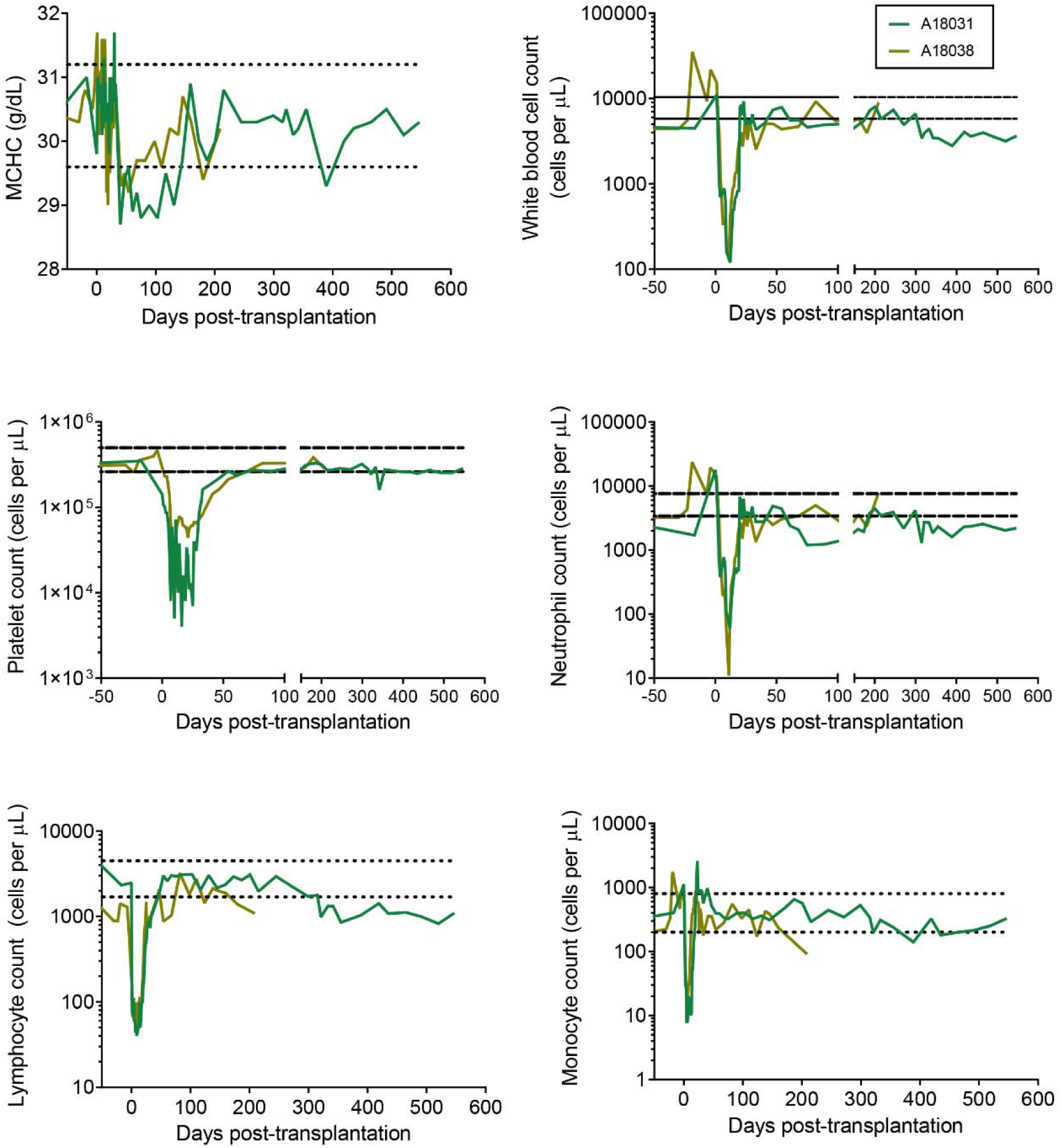
Hematopoietic recovery in both multiplex edited NHP transplants as determined by complete blood cell counts. Dashed lines show normal count range for each lineage.

**Supplemental Fig. 11.**
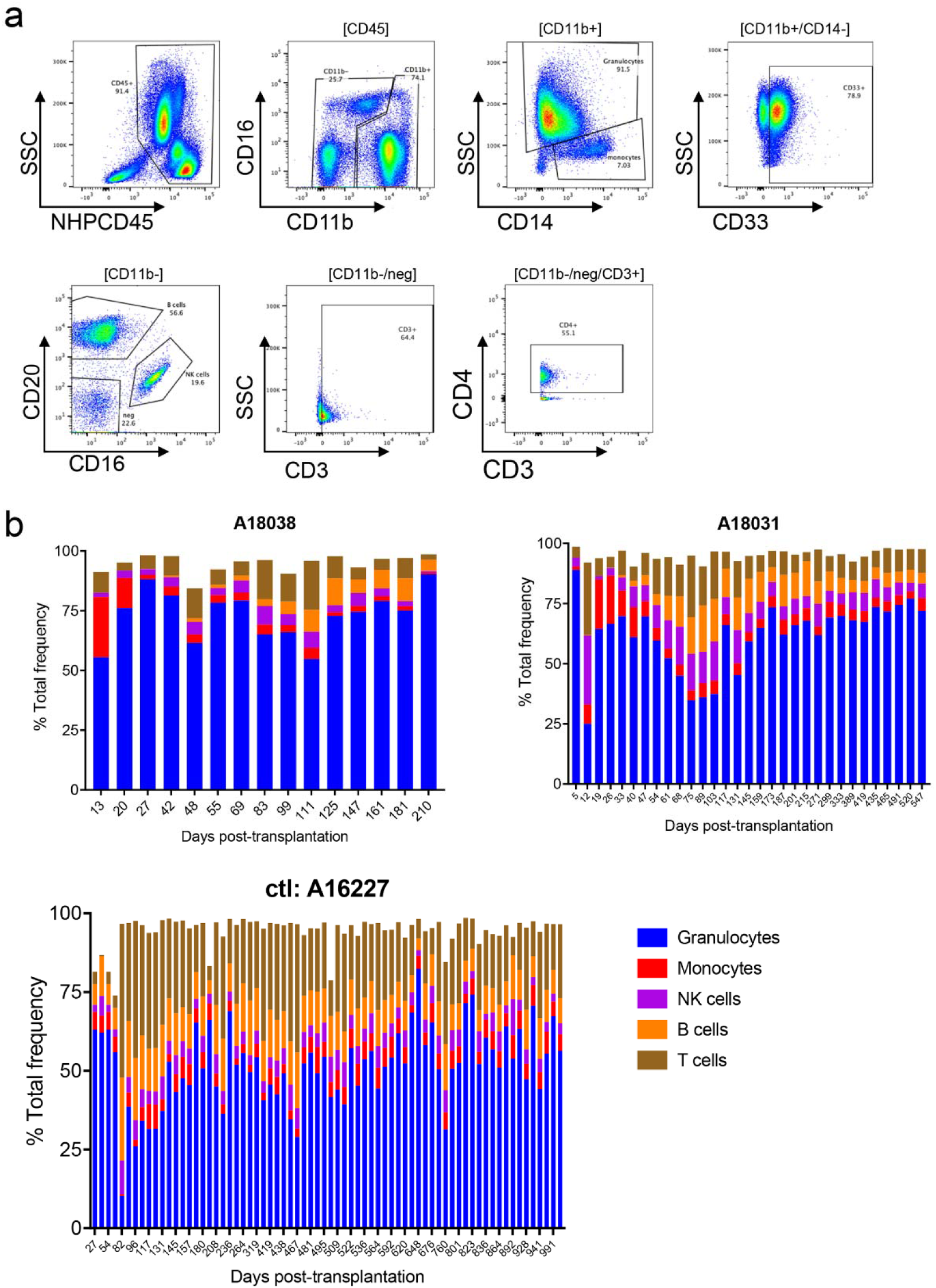
Hematopoietic recovery in both multiplex edited NHP transplants as determined by reconstitution of different blood lineages assessed by flow cytometry. **a,** Representative flow cytometry plots showing gating strategy for blood lineages **b,** Frequency of each blood lineages from the 2 multiplex NHP transplants as compared to an untransplant NHP control (A16227).

**Supplemental Fig. 12.**
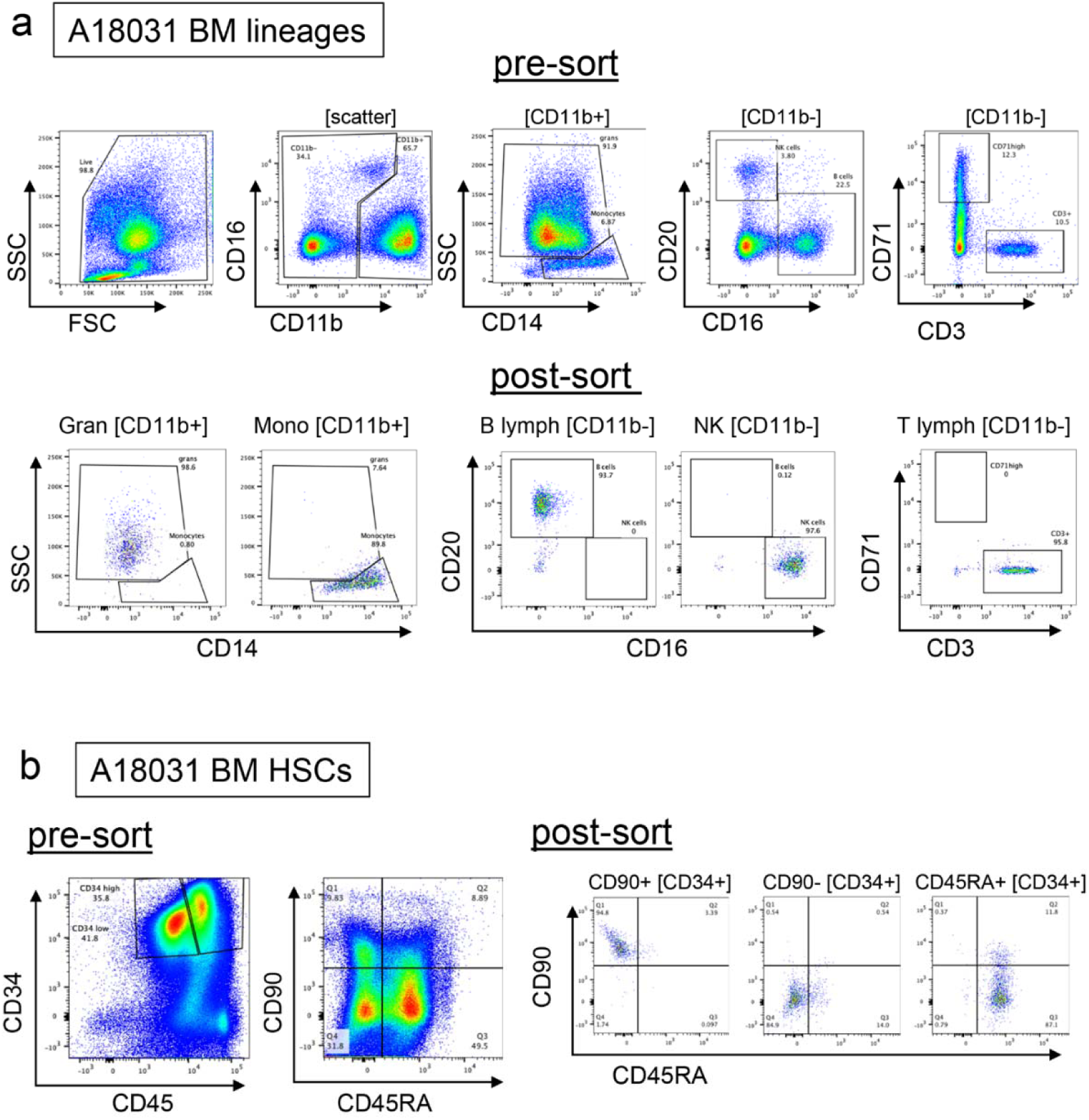
Flow cytometry assessment of bone marrow samples taken from both transplanted animals A18031. **a,** Flow cytometry plots showing gating strategy for blood lineages before (top) and after (bottom) sorting. **b,** Flow cytometry plots showing gating strategy for HSPC subsets before (left) and after (right) sorting.

**Supplemental Fig. 13.**
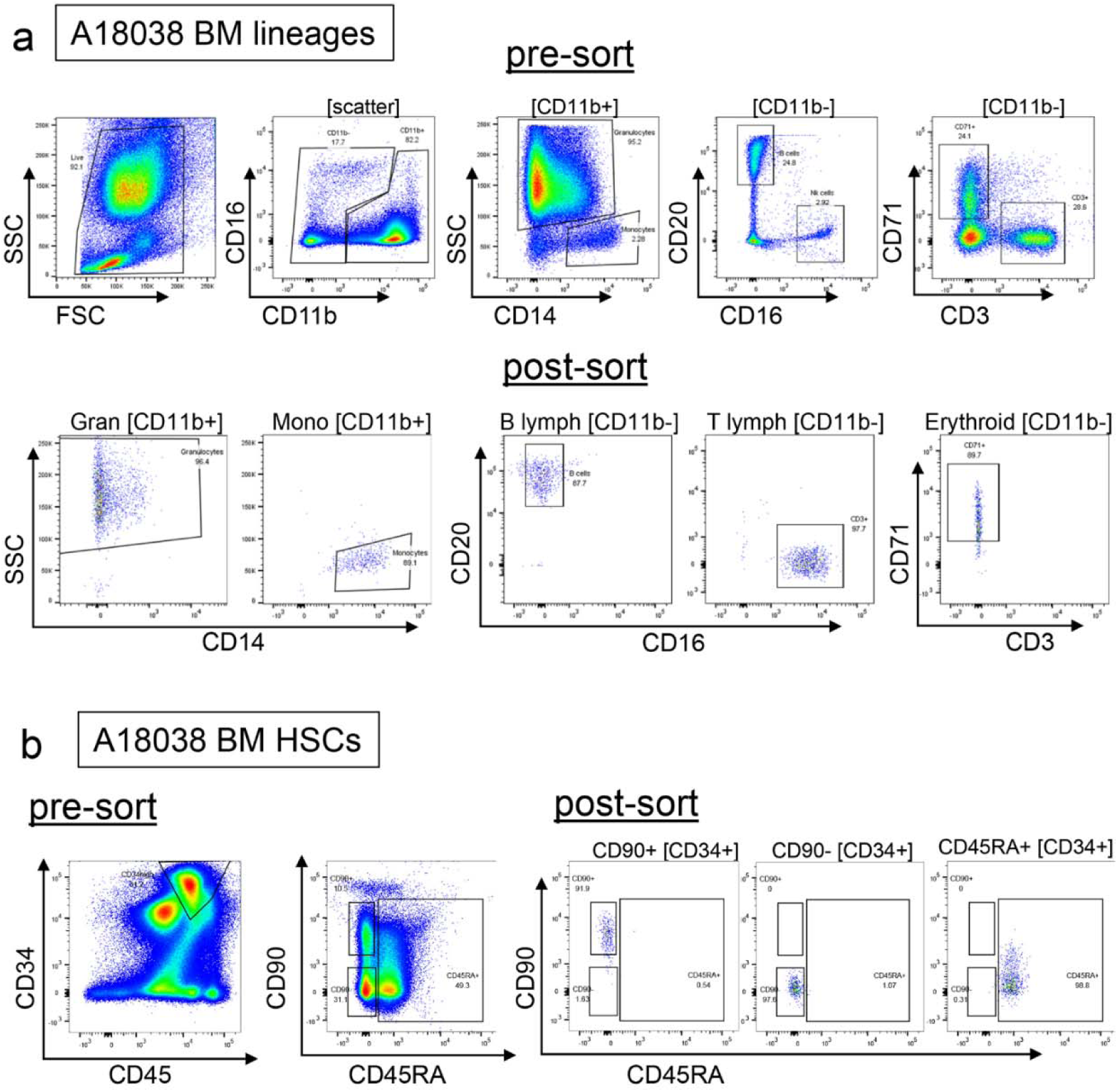
Flow cytometry assessment of bone marrow samples taken from both transplanted animals A18038. **a,** Flow cytometry plots showing gating strategy for blood lineages before (top) and after (bottom) sorting. **b,** Flow cytometry plots showing gating strategy for HSPC subsets before (left) and after (right) sorting.

**Table S1.**
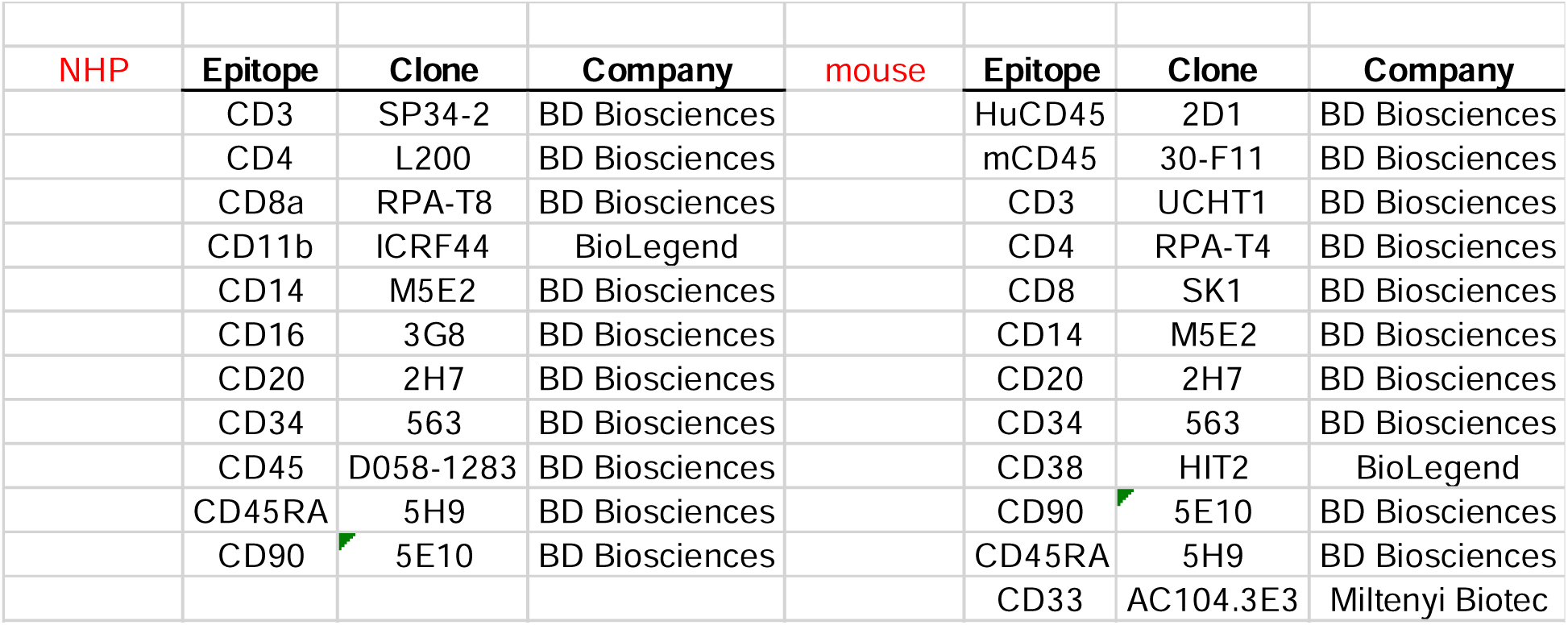
list of antibodies for flow cytometry

**Table S2.**
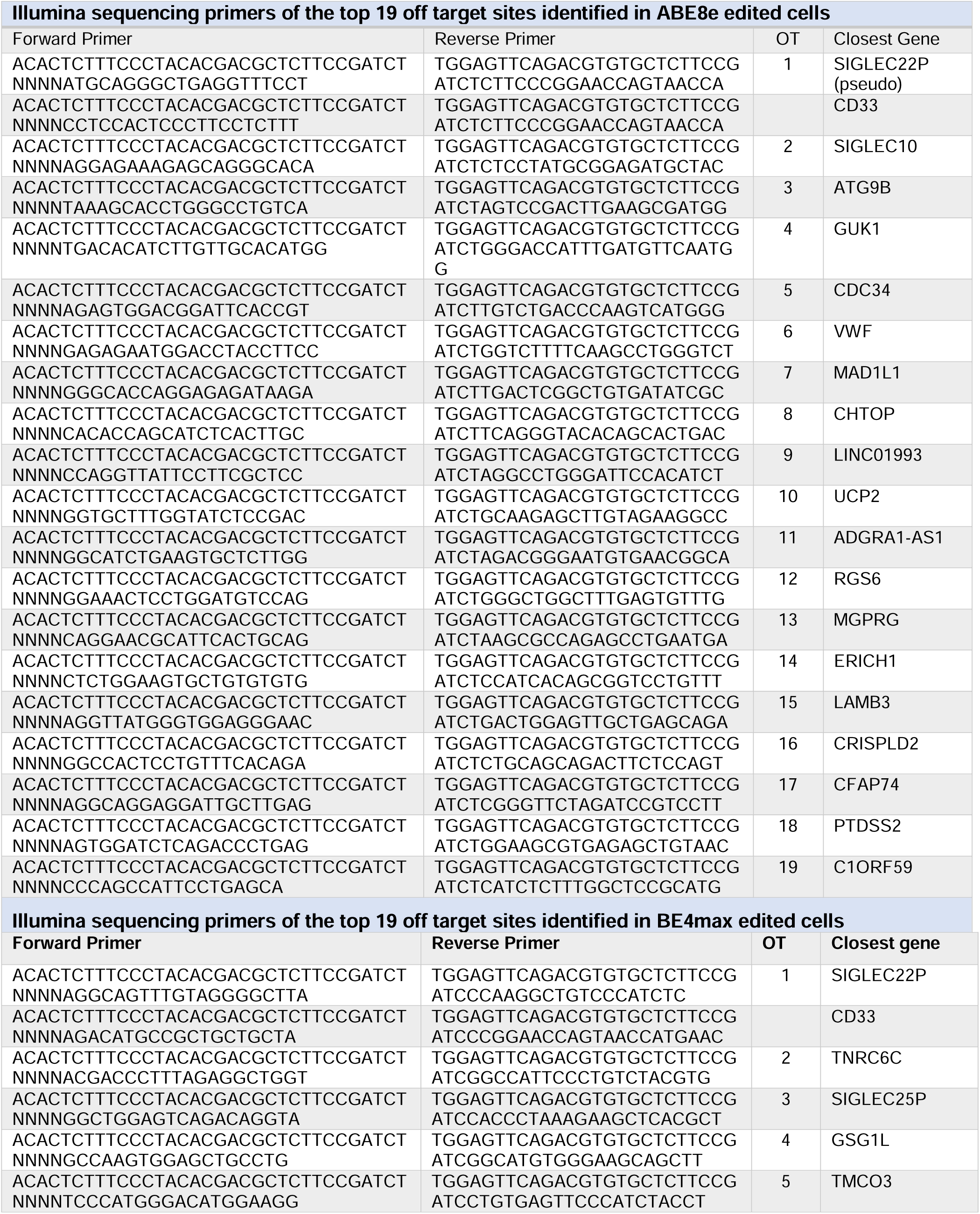
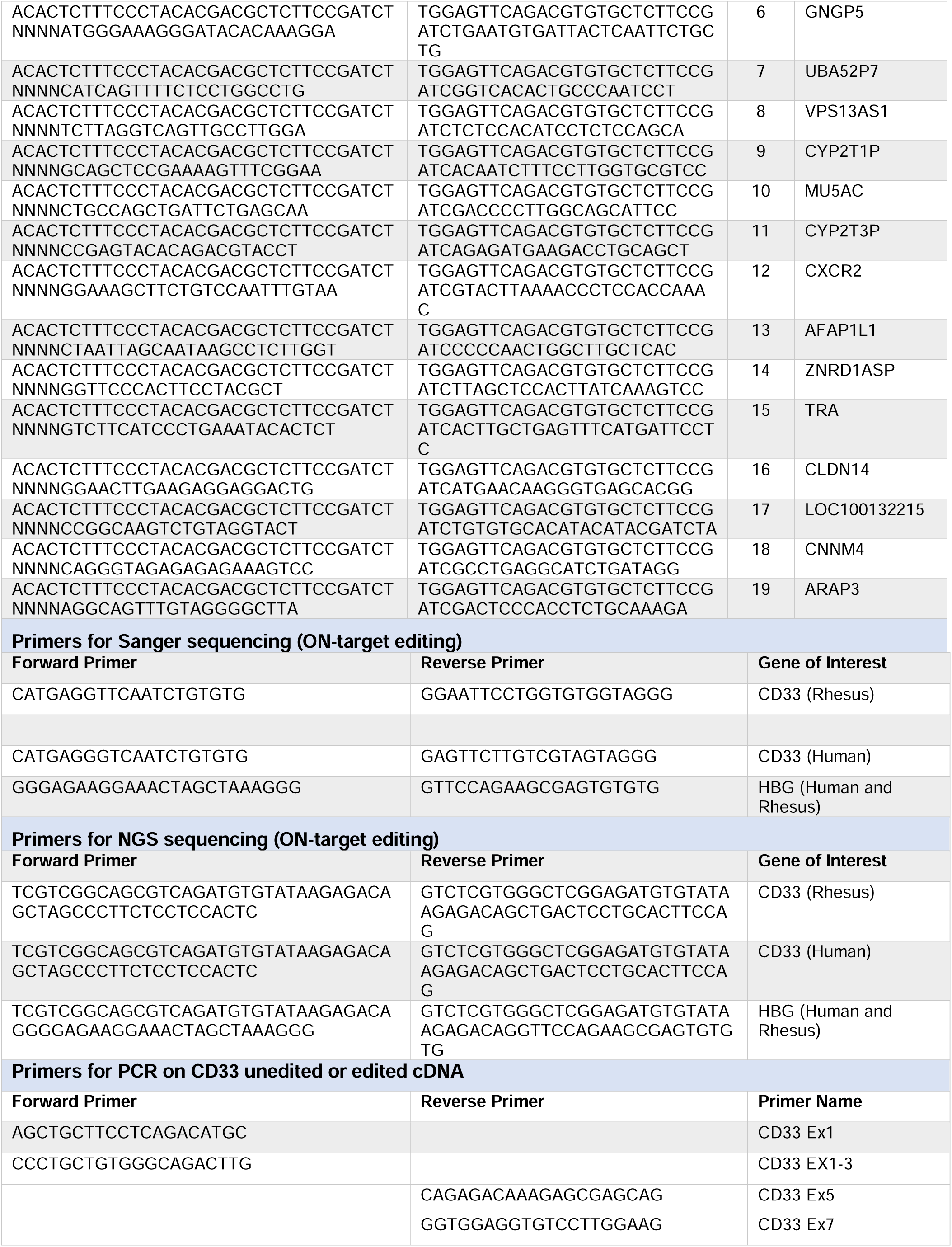
Primer’s list

